# PSICHIC: physicochemical graph neural network for learning protein-ligand interaction fingerprints from sequence data

**DOI:** 10.1101/2023.09.17.558145

**Authors:** Huan Yee Koh, Anh T.N. Nguyen, Shirui Pan, Lauren T. May, Geoffrey I. Webb

## Abstract

In drug discovery, determining the binding affinity and functional effects of small-molecule ligands on proteins is critical. Current computational methods can predict these protein-ligand interaction properties but often lose accuracy without high-resolution protein structures and falter in predicting functional effects. We introduce PSICHIC (PhySIcoCHemICal graph neural network), a framework uniquely incorporating physicochemical constraints to decode interaction fingerprints directly from sequence data alone. This enables PSICHIC to attain first-of-its-kind emergent capabilities in deciphering mechanisms underlying protein-ligand interactions, achieving state-of-the-art accuracy and interpretability. Trained on identical protein-ligand pairs without structural data, PSICHIC matched and even surpassed leading structure-based methods in binding affinity prediction. In a library screening for adenosine A1 receptor agonists, PSICHIC discerned functional effects effectively, ranking the sole novel agonist within the top three. PSICHIC’s interpretable fingerprints identified protein residues and ligand atoms involved in interactions. We foresee PSICHIC reshaping virtual screening and deepening our understanding of protein-ligand interactions.

## Introduction

Interactions between small-molecule ligands and proteins underlie many biological processes. Drugs exert intended effects by selectively interacting with specific proteins. Computational methods for understanding these interactions and predicting associated properties can facilitate cost-effective exploration of vast chemical spaces prior to experimental testing, thereby accelerating drug discovery^1,2^. In response, significant research efforts have been focusing on developing computational methods to accurately predict the binding affinity^3–9^ and functional effects of a small molecule on its protein target^10–13^.

Computational methods generally fall into three approaches based on input requirements. *Sequence-based* approaches use either one-dimensional (1D) input^3^, comprising ligand molecular strings and protein sequences, or two-dimensional (2D) input^4^, featuring ligand molecular graphs and predicted protein contact maps. Although contact maps can be determined through laboratory experiments, computationally predicted versions^14–16^ offer sufficient accuracy^17,18^. *Structure-based* approaches^8,19^ require three-dimensional (3D) protein structures along with 1D or 2D ligand input. 3D protein structures are traditionally derived from experimental sources but increasingly utilizing *in silico* structures, such as those generated by AlphaFold2^20^. However, structure-based methods can be sensitive to errors introduced by *in silico* predictions^21,22^. *Complex-based* approaches^6^ require protein-ligand co-complex structures, differing from structure-based approaches that do not require 3D ligands to be co-bound to protein structures.

Ceteris paribus, a sequence-based approach is preferable because it eliminates the need for enormous cost of experimental structure determination and potential inaccuracies from *in silico* structure prediction. Furthermore, unlike structure-based approaches^10–12^, sequence-based approaches are not sensitive to the protein structural state when predicting the functional effects of a small molecule on its protein target. Instead, sequence-based approaches can directly ascertain binding affinities and the functional effects of protein-ligand interactions^13^. To this end, deep learning methods applied to sequence data offer increasingly accurate predictions^3,23^. However, since deep learning methods fundamentally rely on pattern-matching, deep sequence-based methods can suffer from unrestrained degrees of freedom^24^. This can lead to overfitting to incorrect training patterns, resulting in limited generalizability^9^. Consequently, a substantial performance gap remains between sequence-based methods and both structure-based^8,25^ and complex-based methods^6^.

We argue protein-ligand interaction patterns can be decoded and fingerprinted from sequence data alone. Our core hypothesis, nonetheless, is subject to a crucial condition: deep learning architectures should be endowed with adequate physicochemical constraints to realize this proposition. Compared to previous deep sequence-based methods, this approach could provide a more faithful representation of the underlying protein-ligand interactions, thereby closing the performance gap between sequence-based methods and structure-based or complex-based methods. Recent advancements in structure-based^8^ and complex-based methods^6^ using graph neural networks (GNNs) have alluded to this promising direction. These methods outperform convolutional neural network counterparts by capturing 3D geometric and physicochemical constraints. However, integrating these constraints into sequence-based methods, which operate in 1D or 2D, remains a challenge. Current sequence-based GNNs either use simple concatenation of protein and ligand representations^4,17^, or model interactions using complete pairwise comparisons between every protein residue and ligand atom^9,18^.

Here, we present PSICHIC (pronounced psychic), PhySIcoCHemICal graph neural network, a method designed to adhere to physicochemical principles when decoding protein-ligand interaction fingerprints directly from sequence data. Unlike previous sequence-based models, PSICHIC uniquely incorporates physicochemical constraints to achieve state-of-the-art accuracy and interpretability. When trained on identical protein-ligand pairs without structural data, PSICHIC matched and even surpassed leading structure-based and complex-based methods in binding affinity prediction. PSICHIC also excels in predicting the functional effects of small molecules on protein targets, achieving a high accuracy score of 0.96 in this regard. Intriguingly, PSICHIC’s interpretable fingerprint revealed that the method gained emergent capabilities in deciphering mechanisms underlying protein-ligand interactions directly from sequence data alone. Specifically, PSICHIC identified protein residues in the binding site and ligand atoms involved in interactions, despite being trained on sequence data with only binding affinity labels but without any binding site and residue-atom interaction information. In a proof-of-concept study, we applied PSICHIC to screen a 30,282-compound library for adenosine A_1_ receptor (A_1_R) agonists. Impressively, PSICHIC ranked the only agonist (MIPS1) in the library, according to pharmacological evaluation, within the top 3. MIPS1 exhibits a Tanimoto similarity to the closest known A_1_R agonist of only 0.2, affirming PSICHIC’s potential for novel drug discovery.

## Results

### PSICHIC Framework

PSICHIC presents a framework for learning protein-ligand interaction fingerprints from annotated sequence data (Fig. 1). The input for PSICHIC can be any sequence data comprising a protein-ligand pair with a protein sequence and a ligand’s Simplified Molecular Input Line Entry System (SMILES) string. To train PSICHIC, these pairs need to be annotated with interaction properties like binding affinity and functional effects.

**Fig. 1.**
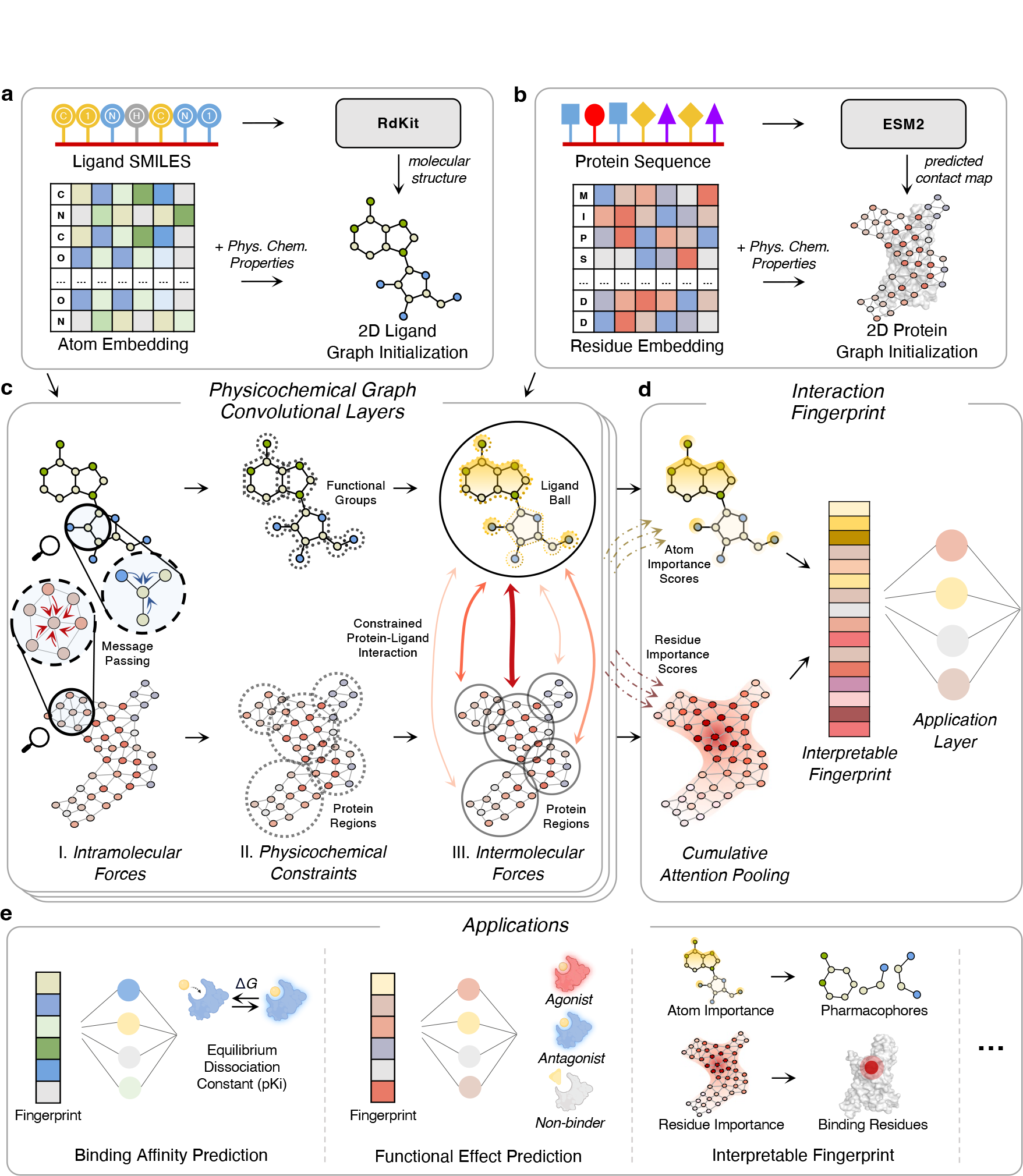
PSICHIC (PhySIcoCHemICal graph neural network). **a**, Ligand SMILES forms an atom graph with atom type embeddings and physicochemical properties, connected by covalent bonds. **b**, Protein sequence forms a residue graph, using ESM2 embeddings and physicochemical properties, connected by predicted contact map from ESM2 protein language model (see Methods). **c**, Over three iterative layers, (I) PSICHIC models the intramolecular forces by passing messages between atoms and between residues using two independent GNNs. (II) PSICHIC imposes physicochemical constraints by grouping ligand atoms into functional groups and protein residues into clustered protein regions. (III) PSICHIC models intermolecular forces in three steps: first, it aggregates ligand functional groups into a ligand ball; second, it calculates interaction strengths between the ligand ball and protein regions; third, PSICHIC disaggregates the ligand ball into updated ligand atoms and ungroups clustered protein regions into updated protein residues to conclude one layer. **d**, After three layers, PSICHIC creates an interaction fingerprint, weighting atoms and residues via importance scores from intermolecular forces. The fingerprint serves as input to a single-hidden-layer network for predictions. **e**, PSICHIC’s interaction fingerprints are generalizable and interpretable across tasks.

PSICHIC initializes the ligand graph from a ligand SMILES (Fig. 1a) and the protein graph from a protein sequence (Fig. 1b). For the ligand, atoms are connected via the covalent bonds to form a molecular graph. Each atom is initialized with an atom type embedding and its physicochemical properties. For the protein, residues are connected via a predicted contact map using ESM2^16^ (see Methods). Each residue is initialized with an ESM2 residue embedding and its physicochemical properties.

PSICHIC models the protein-ligand interaction by interweaving intra- and intermolecular forces through three physico-chemical convolutional layers (Fig. 1c). For each layer, PSICHIC starts modeling intramolecular forces by passing messages between atoms within the ligand, and between residues within the protein using two independent GNNs^26^ (Fig. 1c-I.).

PSICHIC incorporates physicochemical constraints to prepare for intermolecular force modeling (Fig. 1c-II.). PSICHIC groups ligand atoms into functional groups and protein residues into clustered regions. These constraints guide PSICHIC in discovering interaction patterns that adhere to physicochemical principles. For the ligand, the constraints enable recognition of key functional groups for interaction. For the protein, the constraints limit interactions to those between the entire ligand and specific clustered regions of the protein, thereby minimizing the potential to overfit on random, dispersed interactions.

PSICHIC models intermolecular forces through a three-step process (Fig. 1c-III.). First, PSICHIC aggregates ligand functional groups into a single representation, termed a “ligand ball”, after weighing each group’s contribution to the interaction using attentional aggregation (see Methods). Second, PSICHIC employs cross-attention to calculate the interaction strength between the ligand ball and protein regions, identifying probable binding sites. Third, PSICHIC disaggregates the ligand ball into updated ligand atoms and ungroups the clustered protein regions into updated protein residues, completing one cycle of the physicochemical convolutional layer.

PSICHIC runs through three cycles to iterate the intra- and intermolecular forces, during which it accumulates the importance scores for both ligand atoms and protein residues based on their contributions to the intermolecular forces (Fig. 1d). PSICHIC then uses importance scores to summarize the protein and ligand graphs into an interaction fingerprint that captures information about protein-ligand interactions. This fingerprint is directly interpretable, as the scores highlight the residues and atoms crucial for interaction. PSICHIC thereby pinpoints probable protein binding sites and key ligand functional groups. Finally, the application layer utilizes this fingerprint to predict interaction properties. The application of interaction fingerprints extends to both predicting and interpreting interaction properties, including binding affinity and functional effects (Fig. 1e).

### Effectiveness of Interaction Fingerprints

We evaluated PSICHIC’s ability to predict protein-ligand binding affinity using the PDBBind v2016 and v2020 benchmark datasets^27^. The v2016^6,28^ and v2020^8,29^ datasets have standard training and testing splits on approximately 3,000 and 17,000 samples, respectively (see Extended Data Table 1 for dataset statistics). Each sample includes an experimentally determined protein-ligand complex and its binding affinity, expressed in negative log-transformed Ki/Kd values. We compared PSICHIC against leading methods in three categories: complex-based, structure-based and sequence-based. Complex-based methods used complete protein-ligand complex structures. Structure-based methods leveraged experimentally-determined 3D protein structures in the ligand-bound state but lacked 3D ligand information. Sequence-based methods used only ligand SMILES and protein sequences. This comparison ensured that we trained and tested all methods on identical protein-ligand pairs, differing only in information accessibility.

On both the PDBBind v2016 and v2020 datasets, PSICHIC substantially outperformed all other sequence-based methods, achieving an average of 15.4% lower values in prediction errors, RMSE and MAE, as well as 18.2% higher values in correlation coefficient metrics, specifically Pearson’s and Spearman’s correlation coefficients (Table 1). In the case of the PDBBind v2016 dataset, PSICHIC was a close second to the complex-based SIGN, and it outperformed other complex-based methods and all structure-based methods. For the PDBBind v2020 dataset, PSICHIC surpassed all methods in achieving the lowest prediction errors, regardless of whether the methods were sequence-based, structure-based, or complex-based. PSICHIC also closely followed the structure-based TankBind in achieving the best correlation coefficient performance. As indicated in Table 1, complex-based methods generally outperform structure-based methods, which in turn often surpass sequence-based methods. The notable exception to this trend is our sequence-based method, PSICHIC.

**Table 1.**
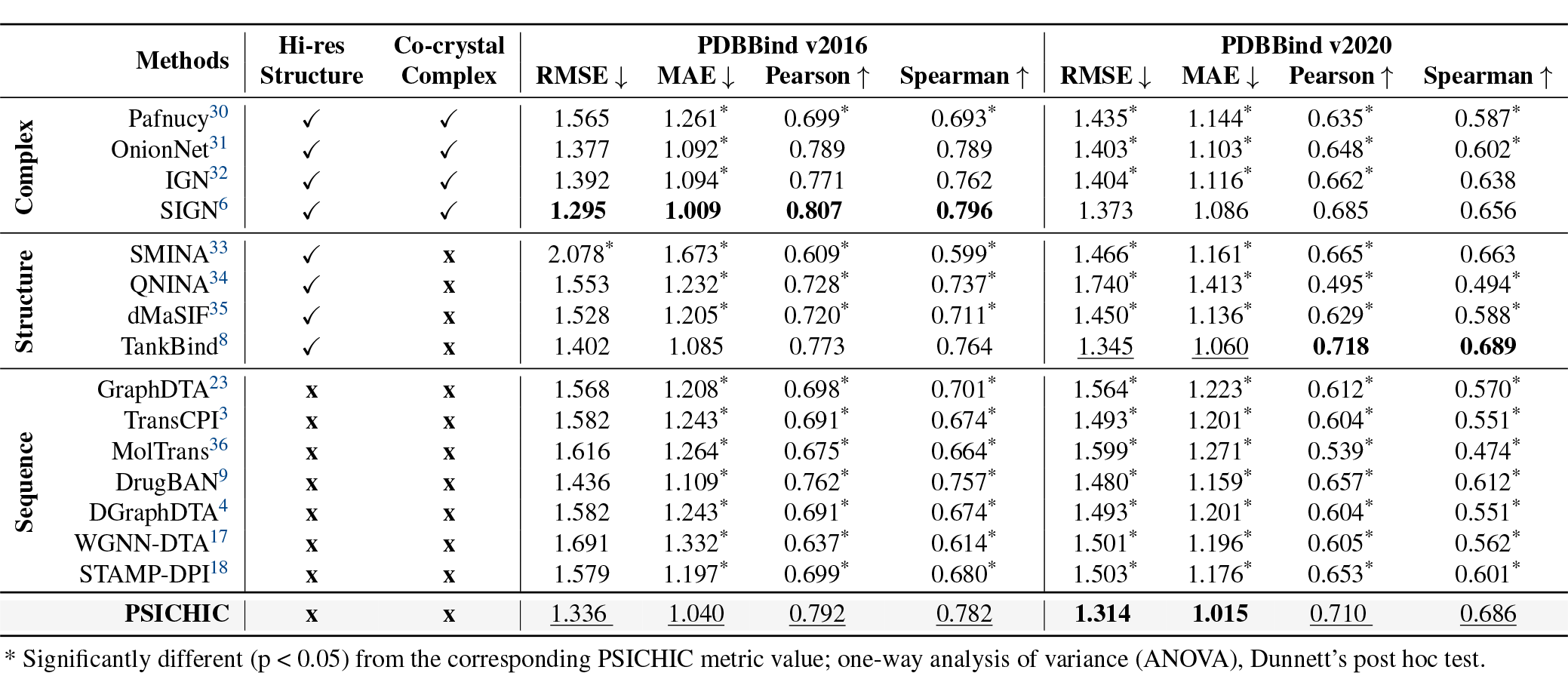
Performance comparison on PDBBind v2016 and v2020 datasets. Prediction errors are measured using root-mean-square error (RMSE) and mean absolute error (MAE), while prediction correlation with experimental affinity values is determined using Pearson’s and Spearman’s correlation coefficients. An upward arrow (↑) denotes higher scores are better, and a downward arrow (↓) denotes the reverse. Average results from five independent runs are reported (see Extended Data Table 2for full statistics). The best performance for each metric is highlighted in **bold**, while the second-best performance is underlined.

Further examining the PDBBind v2020 test set, we observed the leading structure-based, TankBind, and complex-based method, SIGN, experienced higher prediction errors as the numerical value of resolution of the structure increases (Fig. 2a). Note, a larger resolution value, measured in Angstroms, indicates a lower resolution. The rightmost regression plot shows statistically significant relationships for both TankBind and SIGN, with Pearson’s correlation tests yielding p-values less than 0.05. We did not include PDBBind v2016 in comparison because its core test set comprises only high-resolution structures^6,28^.

**Fig. 2.**
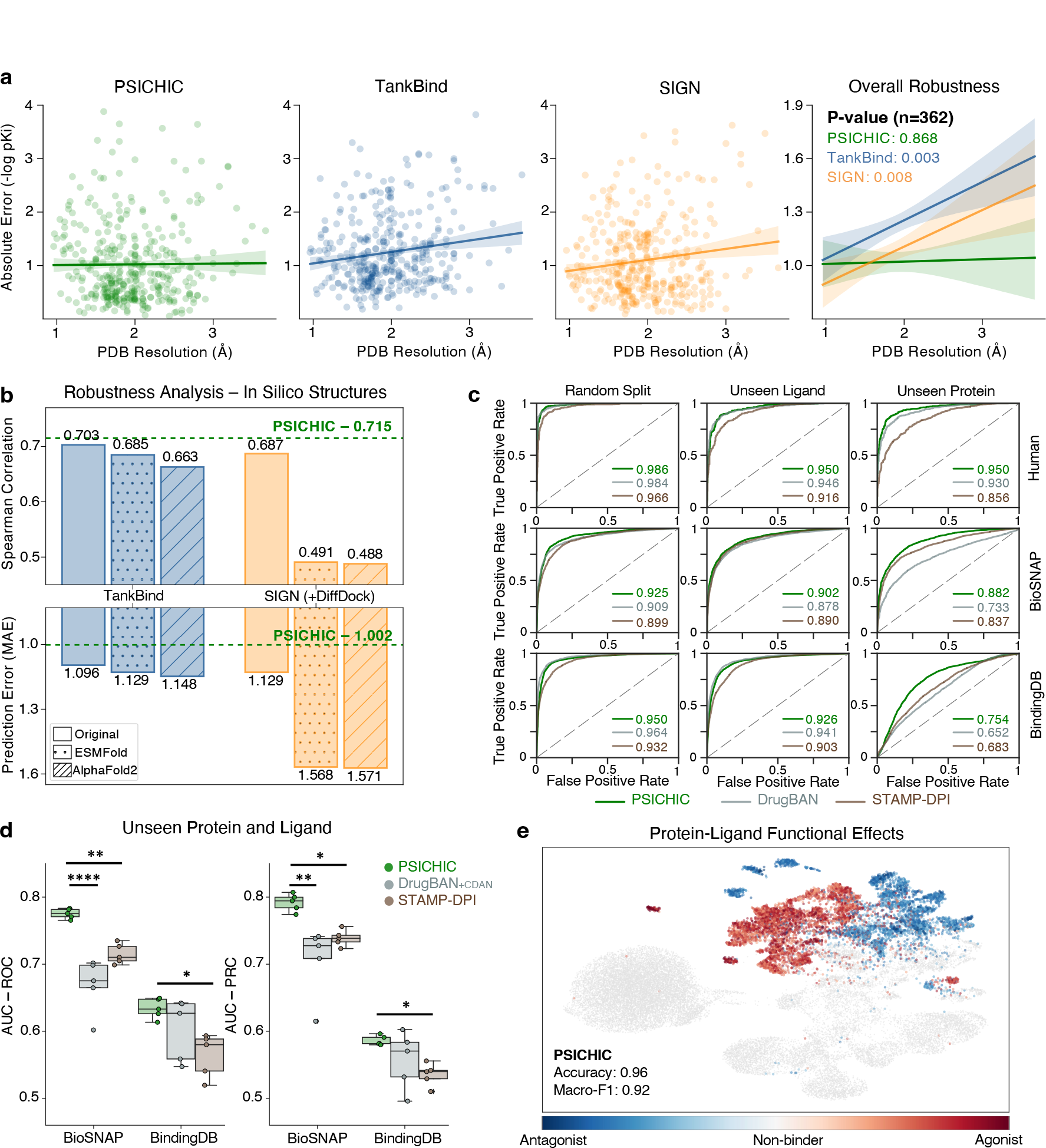
Robustness and Generalizability of Interaction Fingerprints. **a**, Relationship between absolute prediction error and the PDB resolution of protein-ligand complexes in the PDBBind v2020 test set. The rightmost regression plot shows statistically significant linear relationships for TankBind and SIGN, with Pearson’s correlation tests yielding p-values less than 0.05. Error bands represent 95% confidence intervals of linear regression derived from bootstrapping. A larger PDB structure resolution value indicates lower resolution. **b**, Performance comparison of experimentally-determined structures (original PDB structures) versus *in silico* structures (ESMFold^16^ or AlphaFold2^20^) for TankBind and SIGN on PDBBind v2020 test subset. SIGN further used DiffDock^24^ for complex structures. The green line represents PSICHIC’s performance. The subset was chosen due to limitations in processing long sequences by AlphaFold, ESMFold and DiffDock. **c**, ROC curves for best performance of three top-performing sequence-based methods (PSICHIC, DrugBAN and STAMP-DPI) on BioSNAP^37^, BindingDB^38^ and Human^39^ with random split, unseen ligand split and unseen protein split. **d**, Performance comparison for PSICHIC, DrugBAN and STAMP-DPI on unseen protein and ligand splits. AUC-ROC and AUC-PRC are the Area Under the Receiver Operating Characteristics Curve and Area Under the Precision-Recall Curve, respectively. Significance levels with the asterisks indicating the p-values (^*∗*^*P ≤* 0.05,^*∗∗*^*P ≤* 0.01, ^*∗∗∗∗*^*P ≤* 0.0001); ANOVA, Dunnett’s post hoc test. **e**, UMAP of PSICHIC’s fingerprints for agonist, antagonist and non-binding pairs. Color gradient represents experimental potency (negative log-transformed IC_50_/EC_50_). PSICHIC achieved an accuracy of 0.96 and a weighted F1 score of 0.92 on the protein-ligand functional effect test set.

While experimental efforts have elucidated around 200,000 protein structures^40^, these constitute a minuscule proportion of the billions of known protein sequences^20^. In light of this, we evaluated TankBind and SIGN using only PDBBind v2020 sequence data. Since TankBind and SIGN require 3D structures to function, we used AlphaFold2^20^ and ESMFold^16^ to obtain *in silico* 3D protein structures. For the complex-based method SIGN, we further employed DiffDock, the top-performing docking method on PDBBind v2020^24^, to acquire the complex structures. Under this setting, TankBind’s performance only matched the second-best sequence-based method, DrugBAN, while SIGN’s performance deteriorated substantially (Fig. 2b).

Overall, PSICHIC can surpass structure-based and complex-based methods when trained on the same protein-ligand pairs, without requiring costly high-resolution structural or complex information. Furthermore, while leading structure-based and complex-based methods are significantly sensitive to the resolution of structural input, sequence-based PSICHIC’s performance remains unaffected, affirming its robustness.

### Generalizability of Interaction Fingerprints

PSICHIC, relying exclusively on sequence information, has demonstrated an ability to outperform structure-based and complex-based methods. This prompts us to explore its generalizability.

Using sequence-only datasets, Human^36^, BioSNAP^37^, and BindingDB^38^, as benchmark datasets, we applied three splitting strategies: random split, unseen ligand split, and unseen protein split. Each split subdivided the dataset into training, validation, and test sets. The unseen ligand split ensured that ligand scaffolds in the test sets were unseen during model training, while the unseen protein split ensured that protein targets in the test sets were likewise unseen during training. This resulted in nine diverse settings to evaluate model generalizability (see Extended Data Table 1 for dataset statistics). We then compared PSICHIC’s performance against seven other sequence-based methods using binary classification to predict interaction activity. PSICHIC led in performance, as measured by the area under the Receiver Operating Characteristic curve (ROC-AUC), followed by DrugBAN^9^ and STAMP-DPI^18^ (Fig. 2c, Extended Data Fig. 2).

To further assess PSICHIC’s generalizability, we applied a similar dataset splitting strategy to that of DrugBAN^9^, splitting the BioSNAP and BindingDB datasets such that both proteins and ligands in the test set are unseen and substantially different from those in the training set. Like DrugBAN, we excluded the Human dataset due to its insufficient size for effective splitting. When benchmarked against DrugBAN and STAMP-DPI, PSICHIC substantially outperformed both, despite DrugBAN’s use of a specialized Cross-Domain Adaptation Mechanism (CDAN) designed to handle unseen proteins and ligands (Fig. 2d).

In practical virtual screening, while it is important to know the binary interaction activity, determining whether a ligand activates (agonist) or deactivates (antagonist) a protein target is necessary for efficient drug discovery. To address this, we created a protein-ligand functional effects dataset using data from ExCAPE^41^, Papyrus^42^, and Cortellis (Extended Data Fig. 3). Training PSICHIC on this dataset enabled it to predict whether a ligand is an agonist, antagonist, or non-binder for a protein. After training and validating PSICHIC on 70% and 10% of the dataset, respectively, we tested it on the remaining 20%. PSICHIC demonstrated a high accuracy score of 0.96, and a class-equally-weighted F1 score of 0.92 (Fig. 2e).

Overall, PSICHIC has proven its ability to learn generalizable interaction fingerprints, allowing it to accurately predict the binding affinity, interaction activity, and functional effects of small molecules on protein targets. This establishes PSICHIC as a versatile tool in a wide range of drug discovery endeavors.

### Interpretability of Interaction Fingerprints

What insights has PSICHIC gained? Does the reasoning behind PSICHIC rely on comparisons of training instances, or does it grasp the underlying physicochemical principles that dictate binding? To address these questions, we examined the interpretation of PSICHIC on the PDBBind v2020 test set. This set offered co-crystallized structures for evaluation and featured protein-ligand pairs PSICHIC had not encountered during its training.

We examined the relationship between the protein residue proximity to the binding site and the PSICHIC’s assigned residue importance scores. When PSICHIC predicted high binding affinity values, PSICHIC assigned higher importance scores to fewer specific residues, reflecting PSICHIC’s higher confidence in which residues were important for binding. Notably, these residues in fact form the actual binding sites as evidenced by the increase in correlation between residue importance scores and residues’ proximity to the binding site (Fig. 3a). The tendency for the scores to converge on the binding site as predicted affinity values increase is consistent with physicochemical principles: a stronger binding affinity increases the likelihood of a complex system transitioning into a stable, low-energy state.

**Fig. 3.**
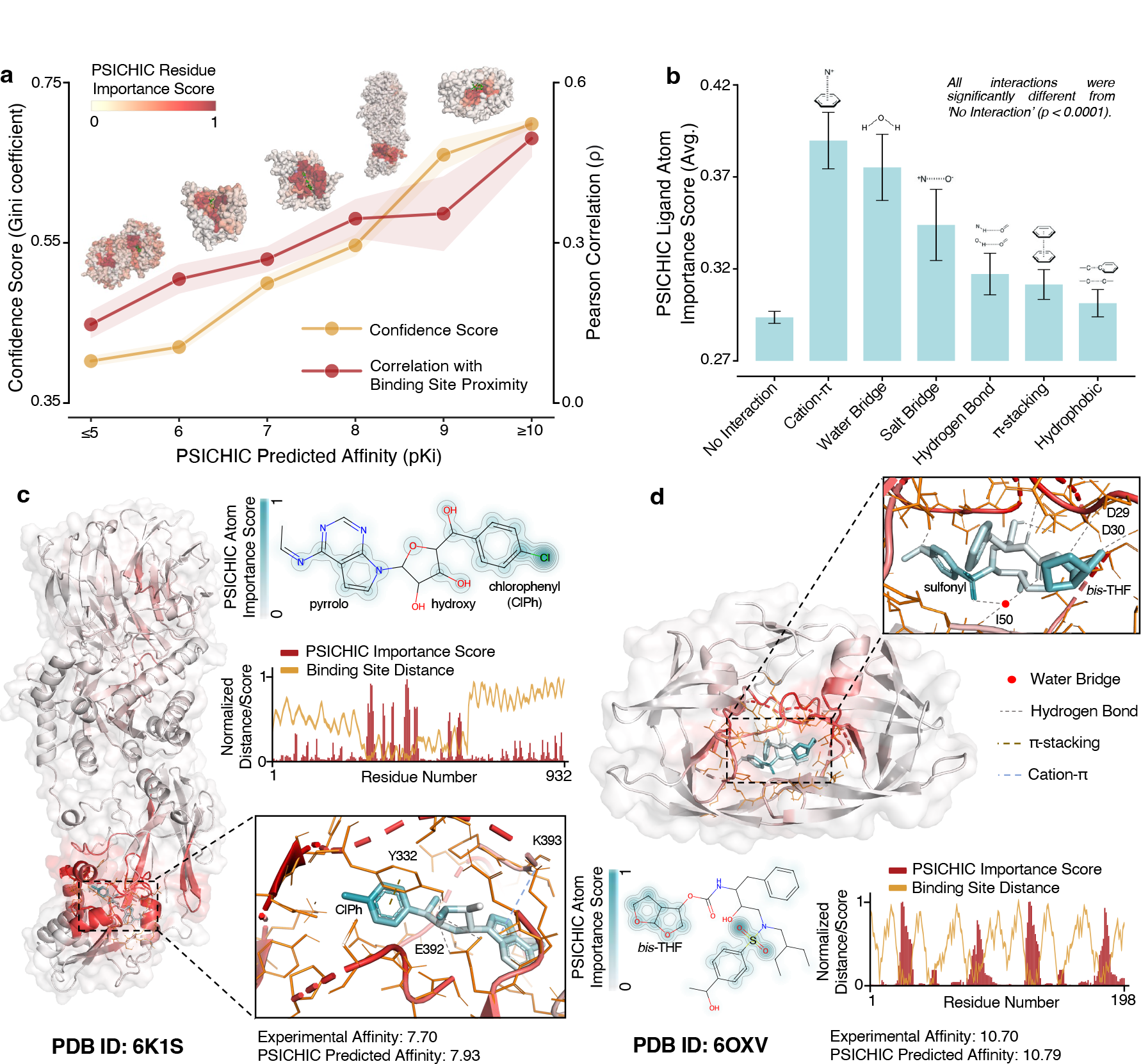
Interpretability of Interaction Fingerprints. **a**, The line plot illustrates relationships based on the predicted binding affinity by PSICHIC: (1) an orange line shows the relationship between affinity (x-axis) and confidence in residue importance scores (y-axis), the latter being measured by the Gini coefficient. (2) a red line depicts the relationship between affinity (x-axis) and the correlation of PSICHIC residue importance scores with residues’ proximity to the binding site (y-axis). The error band (shaded region) of line plots indicates one standard error of the mean (SEM). Five PDB complexes above the line plot are annotated with PSICHIC residue importance scores. These complexes are associated with predicted affinity values of 5.76, 6.66, 7.56, 7.93, and 10.79, respectively. **b**, Bar plot of average PSICHIC ligand atom importance, categorized by interaction type as identified by Protein-Ligand Interaction Profiler (PLIP)^43^ (see Supplementary Methods 7). Error bars denote SEM. The category “No Interaction” denotes atoms that do not form any interactions. All interaction types significantly differ from “No Interaction” (p < 0.0001); ANOVA, Dunnett’s post hoc test. Interpretation by PSICHIC for **c**, PDB 6K1S, and **d**, PDB 6OXV. Atom and residue importance scores are projected back onto the complex structure. The ligand plot highlights atoms that PSICHIC attributes significant weight to, with insets illustrating the interactions these atoms form with the binding residues. The bar-line plot and protein structure demonstrate higher importance scores for residues closer to the binding site.

To assess the interpretability of PSICHIC on the ligand side, we annotated each ligand atom based on the interaction type it formed with binding residues. Comparing the average importance scores assigned by PSICHIC to these atoms, we showed that atoms forming interactions received significantly higher scores than non-interacting atoms (Fig. 3b). While each atom within a ligand contributes to its function, the ability of PSICHIC to emphasize interacting atoms makes it a valuable tool for pharmacophore analysis (see Extended Data Fig. 4 and Supplementary Methods 8).

As case studies, we examined two high-affinity PDB complex structures: 6K1S and 6OXV. For 6K1S, an arginine N-methyltransferase 5-inhibitor complex^44^, PSICHIC highlighted residues around the binding site and identified critical ligand functional groups, namely chlorophenyl, pyrrolo, and hydroxyl groups, which interact with Y332, K393, and E392, respectively (Fig. 3c). In PDB 6OXV, an HIV-1 protease-ligand complex^45^, PSICHIC also assigned significant weights to binding residues (Fig. 3d). PSICHIC emphasized the ligand’s bis-tetrahydrofuran moiety, a key feature aiding in resistance evasion^46^, and the sulfonyl group that forms a water bridge with I50. These results support our core hypothesis that imposing physicochemical constraints allows interaction patterns to be decoded and fingerprinted solely from sequence data. This claim is further corroborated by PSICHIC’s failure to identify binding residues when these constraints are removed (Extended Data Fig. 5).

PSICHIC’s emergent ability to decode the fundamental physicochemical mechanism underlying protein-ligand interaction is unique. It accomplishes this by training exclusively on sequence data with binding affinity labels, without requiring any information about binding sites and residue-atom interactions. To the best of our knowledge, this is the first of its kind.

### Virtual Screening with Interaction Fingerprints

Having established PSICHIC’s efficacy and interpretability, we proceeded to assess its practical application on virtual screening to identify A_1_R agonists, an important G protein-coupled receptor implicated in cardiovascular and neuronal diseases^47^. PSICHIC was deployed to virtually screen an in-house MIPS library of 30,282 diverse compounds, with low Tanimoto similarity to known A_1_R agonists. Comparing the MIPS library with existing datasets, only 4.3% of the library compounds have a Tanimoto Similarity greater than 0.5 with any A_1_R bioactivity data in ChEMBL. This proportion dips to a mere 0.6% for the PDBBind v2020 dataset. Methods trained on these databases would thus need exceptional extrapolative abilities to effectively screen the MIPS library. To address this, we collated large-scale sequence databases, ExCAPE and Papyrus, with our functional effect dataset, creating a composite set of 3 million samples annotated with binding affinity and functional effect labels (Extended Data Fig. 6). Notably, 94.3% of MIPS library compounds had a Tanimoto score above 0.5 with this dataset. We then trained PSICHIC on this dataset via a multi-task optimization and diverse interaction sampling approach (Methods, Extended Data Fig. 7, 8). We note that model optimization on such a vast and diverse dataset could only be achieved using sequence-only information or by augmenting the dataset with *in silico* structures^16,20^.

After training, PSICHIC ranked the MIPS library compounds based on predicted binding affinities and functional effects (Fig. 4a). We assessed the efficacy of this ranking against experimental data from a high-throughput screening campaign of the MIPS library targeting human A_1_R (Fig. 4a, b). This pharmacological evaluation involved two steps: initial high-throughput screening of the MIPS library to assess whether the molecules bind to A_1_R, followed by pharmacological characterization to determine the ability of binding compounds to activate the receptors. The initial high-throughput screening identified three ligands (MIPS1, MIPS2, and MIPS3) that bind to human A_1_R. Among these three, only MIPS1 induced an inhibition of cAMP accumulation that was comparable to the effect of the agonist, NECA, and showed no effect on non-transfected cells (see Methods and Fig. 4b), suggesting that it is the sole agonist in the MIPS library.

**Fig. 4.**
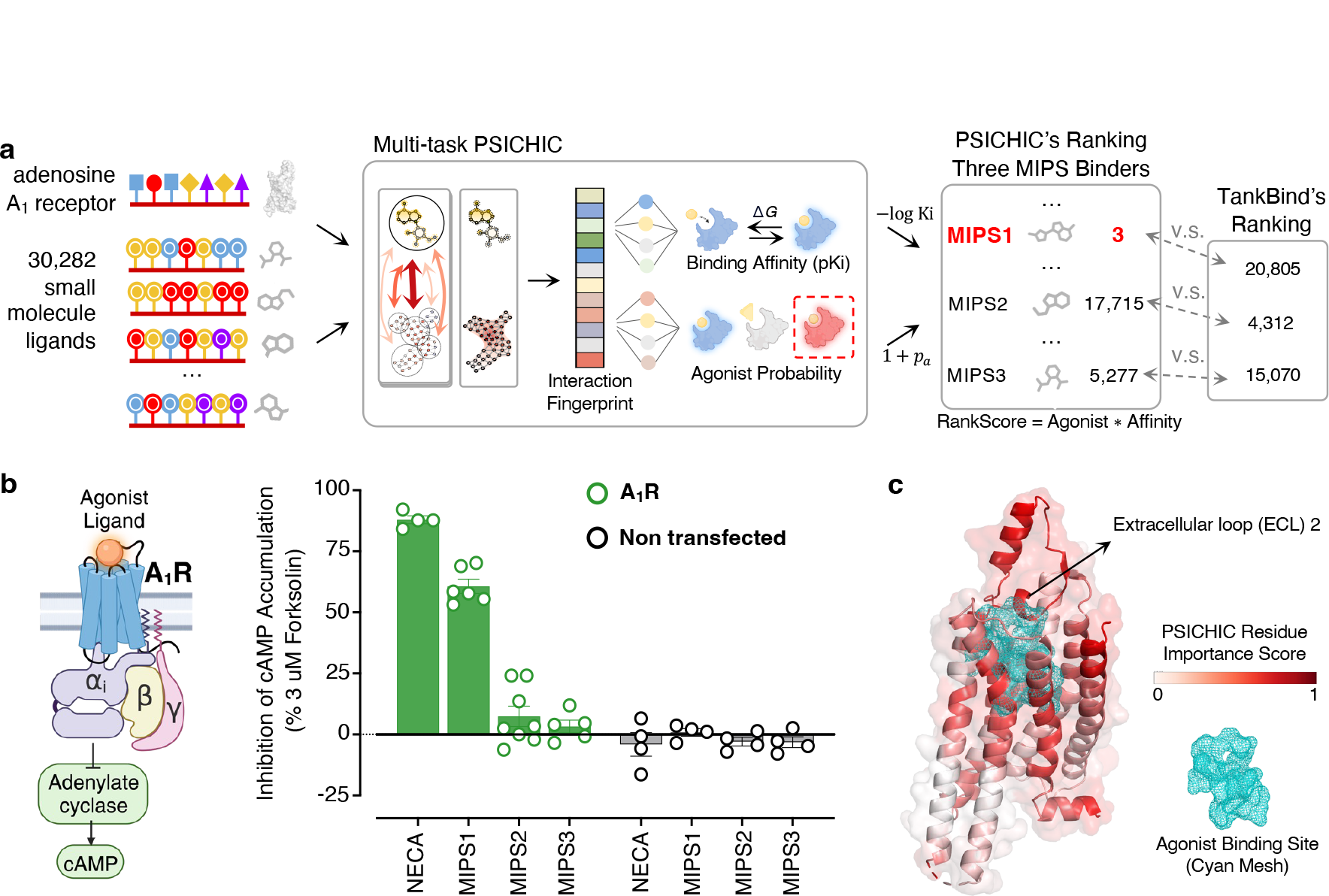
Virtual Screening with Interaction Fingerprints. **a**, PSICHIC-based *in silico* screening of our in-house MIPS library of 30,282 compounds to identify potential A_1_R agonists. PSICHIC, trained on a comprehensive database, combined both binding affinity and functional effect predictions to rank compounds (see Methods and Extended Data Fig. 7, 8). Compounds were scored by multiplying binding affinity (pKi) with (1 + agonist probability). This ranking was validated experimentally against high-throughput screening data for the same library. This screen identified three ligands (Binders: MIPS1, MIPS2, and MIPS3) that bind to human A_1_R. PSICHIC ranked these binders at 3, 17,715, and 5,277, whereas TankBind ranked them at 20,805, 4,312, and 15,070 respectively. **b**, Pharmacological characterization to determine the ability of the three compounds to activate the receptors using cAMP accumulation assay in FlpIN CHO cells stably expressing human A_1_R. MIPS1 induced a robust inhibition of cAMP accumulation comparable to NECA, an A_1_R agonist. Among 30,282 ligands, PSICHIC ranked the only agonist (MIPS1) in the library in the top three, landing it in the highest 0.01 percentile. **c**, Residue Importance Score from PSICHIC’s modeling showed that key residues in A_1_R-MIPS1 interaction are primarily located in the extracellular area above the orthosteric agonist site. A_1_R structure was sourced from PDB: 6D9H^47^, and the cyan mesh represents the adenosine agonist binding site.

PSICHIC’s predictive ranking impressively placed the only agonist compound, MIPS1, in the top 3, or the upper 0.01 percentile (Fig. 4a). This is despite MIPS1’s low 0.2 Tanimoto similarity to any known A_1_R agonists in both the training data and the ChEMBL database. In contrast, the leading structure-based method, TankBind, ranked MIPS1 at 20,804. PSICHIC also assigned low ranks to two other compounds unlikely to be agonists (Fig. 4b). Examining PSICHIC’s interpretation of the A_1_R-MIPS1 interaction, it prioritized residues in the A_1_R orthosteric agonist site and extracellular loop 2, key regions known to significantly influence agonist binding and function^48^. Overall, this validates PSICHIC as a powerful tool for discovering novel small-molecule drugs, and advancing our understanding of protein-ligand interactions directly from sequence data.

While the focus has been on the virtual screening of A_1_R agonists, PSICHIC is designed for application beyond a single protein target. Through multiple benchmark evaluations across both binding affinity and functional effect tasks presented in this paper, PSICHIC has also proven its ability to accurately predict properties of a broad spectrum of protein-ligand interactions, including those involving novel proteins and ligands. Bearing this potential in mind, we have released a user-friendly, open-source online platform of PSICHIC’s virtual screening application, integrated with Google Colaboratory for easy web-based interaction (see Supplementary Methods 12); the code is also available on GitHub for local deployment.

## Discussion

A protein-ligand interaction fingerprint describes the features of specific interactions occurring between a ligand and the binding pocket residues of a protein. Traditionally, these fingerprints are derived from 3D protein-ligand complexes^49^, a costly process shown here to be sensitive to input resolution quality. In contrast, PSICHIC leverages only sequence data, offering a distinct approach for obtaining interpretable protein-ligand interaction fingerprints. By incorporating constraints, PSICHIC exhibits emergent learning of physicochemical mechanisms, allowing it to unveil complex interaction patterns and to match and even surpass structure-based and complex-based methods. PSICHIC eliminates the need for high-resolution 3D data and paves the way for robust learning on large-scale sequence databases.

As a proof-of-concept, we demonstrate that PSICHIC is effective across diverse drug discovery scenarios and can reliably screen potential drug candidates, including those with low similarity to known compounds. The interpretation of fingerprints from sequence data can facilitate pharmacophore analysis and provide binding site information at a negligible cost. Requiring only sequence data for operation, PSICHIC could serve as a universally useful tool in drug discovery. We also anticipate its utility in the design of new small-molecule therapeutics. Future de novo ligand design programs could integrate PSICHIC’s fingerprints that are inherently interpretable to optimize molecular structures.

We have made our data, code, and optimized model available to the broader scientific community. With proven robustness and efficacy across applications, PSICHIC holds promising potential for future developments and stands poised to meaningfully impact the landscape of virtual compound screening and the design of innovative small molecule therapeutics.

## Methods

### PSICHIC Architecture

#### Graph Network Initialization

PSICHIC first preprocesses 1D protein sequence and ligand SMILES strings into a pair of 2D protein and ligand graphs.

For a given 1D amino-acid (AA) protein sequence with *N*_P_ number of residues, PSICHIC generates the 2D protein contact map graph with *N*_P_ residue nodes. The protein graph is characterized by **P**^(0)^, **A**_P_, and 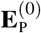, representing the residue node features, the contact map matrix which connects the nodes, and the edge features describing these connections, respectively. To lift the 1D sequence to 2D graph, PSICHIC uses the protein language model ESM2^15,16^, as described in WGNN-DTA^17^, to obtain the predicted contact map matrix, **A**_P_. Edge features, 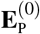, are obtained using a Gaussian radial basis function on predicted contact probabilities from ESM2. The residue node features encode the residue type feature and physicochemical properties. Without extra computation, the residue type feature is obtained from the final layer embeddings of ESM2, 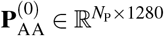, when predicting the contact map. The physicochemical properties, 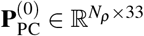, encode the residue weights, pK values, hydrophobicity, and whether the residue is aliphatic, aromatic, neutral, or charged. Both features are transformed and summed using two multilayer perceptrons (MLPs) into the input residue features, 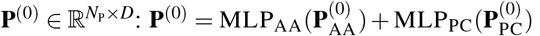, where *D* represents the number of hidden dimensions of PSICHIC network configuration (see PSICHIC Optimization below).

For a given 1D ligand SMILES string with *N*_L_ number of atoms, PSICHIC acquires the 2D molecular graph with *N*_L_ ligand atoms. The ligand molecular graph is described by **L**^(0)^, **A**_L_, and 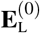 representing the atom node features, the covalent bond matrix which connects the atom nodes, and bond edge features, respectively. To lift the 1D smiles to a 2D graph, PSICHIC uses RdKit to extract the bond information that connects the atoms, **A**_L_, and the bond types. The types of bonds (single, double, triple, aromatic) are transformed into the bond features, 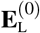, using a learnable embedding matrix. Atom node features encode the atom type feature and physicochemical properties. The atom type features,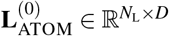, are obtained based on one of the 9 possible element groups (B, C, N, O, P, S, Se, Halogens, and Metals). The physicochemical properties feature,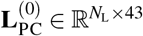, includes the atom degree, the number of implicit Hs, the atom hybridization, and whether it is a chiral atom. The ligand atom input features,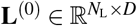, are obtained by 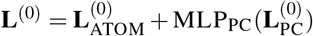.

#### Physicochemical Graph Convolutional (PGC) Layer

In each layer, PSICHIC considers the protein graph, **P**^*l*^,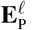, **A**_P_, and the ligand graph, **L**^*l*^,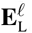, **A**_L_, as inputs to model intramolecular and intermolecular forces. After each layer, PSICHIC acquires the updated protein residue features, **P**^*l*+1^, and ligand atom features, **L**^*l*+1^, along with the importance scores for the protein residues,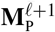, and for the ligand atoms,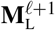. These scores illuminate the significance of the protein residues and ligand atoms in the formation of protein-ligand interactions. In general terms, the operation of each layer can be expressed as:

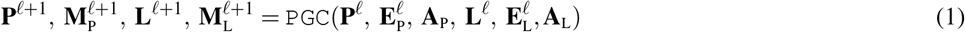

where the PGC layer operates by independently modeling intramolecular forces (Eq. 2), injecting physiochemical constraints (Eq. 3), and, deducing the protein-ligand intermolecular forces (Eq. 4):

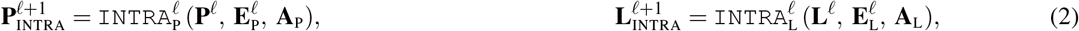

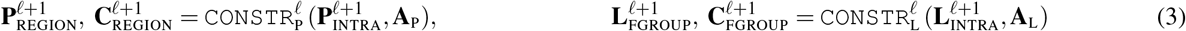

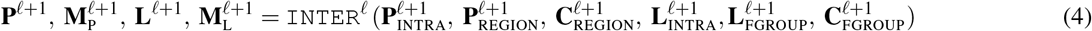

Briefly, the intramolecular forces operation (Eq. 2) includes the use of two independent graph neural networks (GNNs),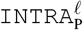 and 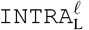, which facilitate the passing “messages” between residues in the protein graph and between atoms in the ligand graph. Following the intramolecular forces operation, the protein residues,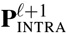, and ligand atoms,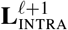, are enriched with information of their neighboring counterparts. The physicochemical constraints operation (Eq. 3) then employs the residue contact map, **A**_P_, and atom bond connection, **A**_L_, to calculate the group assignment matrix for protein residues,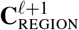, and ligand atoms,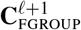. These assignment matrices are subsequently used to cluster protein residues into protein regions, represented as 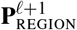, and to organize ligand atoms into their respective functional groups,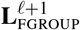. The operation of intermolecular forces (Eq. 4) first computes attention maps as functional group importance scores,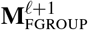, to characterize the significance of functional groups in the protein-ligand interaction. Using 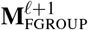, PSICHIC aggregates ligand functional groups into a single ligand representation, termed a “ligand ball”. The interaction strengths are then deduced between the ligand ball and various protein regions, as described by the corresponding importance scores,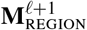, that represent the propensity of specific protein regions for protein-ligand interaction. Leveraging these region importance scores,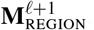, PSICHIC propagates the features from the protein regions to the ligand, and vice-versa. Finally, PSICHIC unpools protein regions to updated protein residues, **P**^*l*+1^, and the region importance scores to residue importance scores,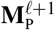, using the region assignment matrix,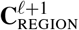. Similarly, PSICHIC maps the functional groups back to ligand atoms, **L**^*l*+1^, and functional group importance scores to atom importance scores,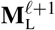, using the functional group assignment matrix,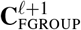.

##### Modeling Intramolecular Forces

PSICHIC utilizes Principal Neighbourhood Aggregation (PNA)^26^ as the GNN backbone to model intramolecular forces. PNA uses four aggregators and three degree-scalers to model the message passing algorithm between residues in the protein graph, as well as between atoms in the ligand graph. This enables the capture of essential physicochemical properties critical for the formation of protein-ligand interactions^50^, including the statistics of aromaticity and hydrogen bond acceptor of neighboring protein residues or ligand atoms. In contrast to the original implementation, PSICHIC substitutes the attenuation scaler with a linear amplification scaler to amplify signals from dense protein regions, a crucial factor given that protein binding sites are frequently characterized by residue density or “buriedness”^51^. A residual connection is added following the PNA message-passing operation, allowing the intramolecular operation to be expressed as follows:

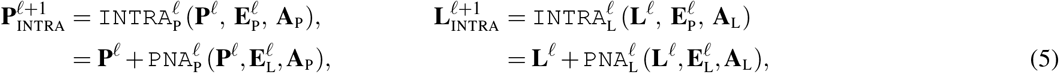

##### Injecting Physicochemical Constraints

To ensure the patterns of protein-ligand interactions discovered through optimization of PSICHIC during training are physicochemically sound, PSICHIC injects constraints by grouping protein residues into protein regions,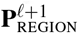 by using the cluster assignment matrix,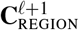, and grouping ligand atoms into functional groups,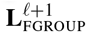, by using the atom assignment matrix,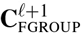:

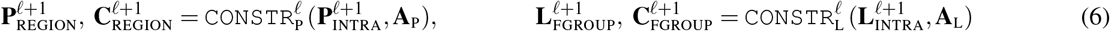

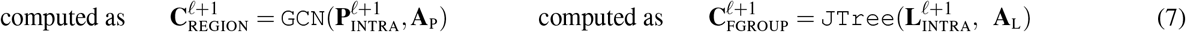

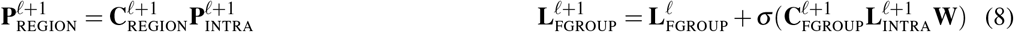

For the protein, the residues that are closely in contact are assigned into clusters by optimizing a continuous clus-ter assignment matrix,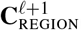, using a graph convolutional network (GCN). The GCN is mathematically defined as 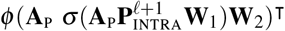 where *σ* is a ReLU function, *φ* is a softmax function, **W**_1_ *∈* ℝ^*D×*2*D*^ and **W**_2_ *∈* ℝ^2*D×K*^ are the parameters of GCN that are trained on the minCUT unsupervised objective, **O**_MINCUT_52 (see Application Layers and Objective Functions below). The *K* dimension in **W**_2_ represents the hyperparameter to set the number of assigned clusters. We obtain *K* number of protein regions with features of *D* dimensions,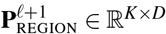, by pooling residues features based on the cluster-assigned residues.

For the ligand, atoms that belong to the same functional groups are likely to have the same importance. Hence, PSICHIC assigns the atoms to their functional groups, which are represented as “structural cliques” identified by junction tree decomposition algorithm^53,54^. The clique assignments,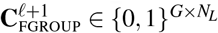, are pre-computed efficiently so that ligand atom features can be assigned into *G* number of functional groups with features of *D* dimensions,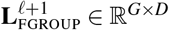. Since the assignments are fixed across different layers, PSICHIC adds features of functional groups from previous layers,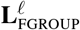, as residuals. For the first layer,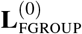 is obtained from a learnable embedding matrix based on whether the functional group is a singleton, ring, bond or bridged compound.

##### Modeling Intermolecular Forces

PSICHIC calculates the intermolecular forces in three steps.

First, PSICHIC consolidates the ligand functional groups into a single ligand representation (visually akin to a ligand ball). This is accomplished by calculating attention map scores,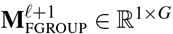, for each functional group. These scores characterize potential significance of the functional groups in the protein-ligand interaction, and PSICHIC aggregates the functional groups into a ligand ball based on these scores:

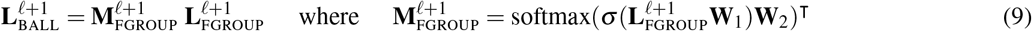

Secondly, PSICHIC deduces the interaction strength between the ligand ball and protein regions to identify the favorable binding site. This is encapsulated in the region importance scores,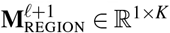. This process is akin to treating the ligand feature as a query or “probe” to find the protein regions that have the highest similarity, as indicated by dot product:

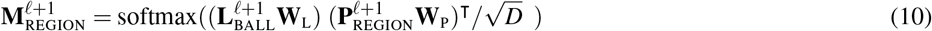

Thirdly, PSICHIC exchanges features between the protein and the ligand based on the favorability of different regions,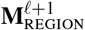:

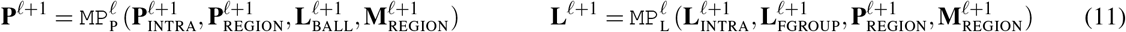

where the update operation is computed for the protein and ligand respectively as:

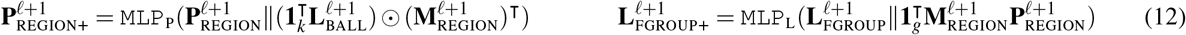

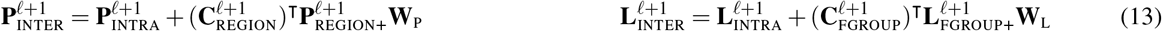

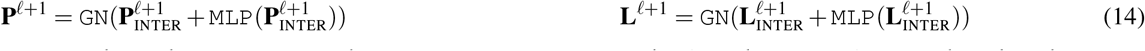

Eq. (12) represents the exchange of features between *K* protein regions and *G* ligand functional groups, based on the region importance scores,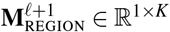.

For the protein regions, the ligand ball features,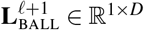, are broadcasted and repeated,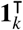, to match the *K* protein regions,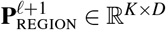. PSICHIC then performs an element-wise multiplication between the broadcasted ligand feature and the transposed region importance scores,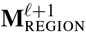. Each row of the element-wise multiplication assigns a weight to *K* repeated ligand ball features, based on the strength of the interaction between different regions. This way, the ligand feature is passed on to protein regions that are deemed more favorable for binding. The weighted ligand features are concatenated,∥, with the region features and fed into an MLP for protein region feature updates.

For the ligand functional groups, protein region features,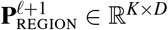, are weighted and pooled using the region importance scores,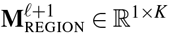. This way, features of protein regions that are more important for interaction will be passed to the ligand more heavily. The pooled protein features are then broadcasted and repeated,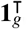, to match *G* ligand functional groups. The weighted protein features are concatenated,∥, with the ligand functional group features and fed into an MLP for ligand functional group feature updates.

Eq. (13) unpools the protein regions and ligand functional groups back to their residue and atom forms to acquire the updated protein residues,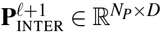, and ligand atoms,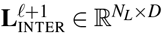. Eq. (14) concludes the PGC layer by updating protein residues and ligand atoms using MLP and Graph Normalization (GN)^55^ operations. Lastly, PSICHIC also unspools the protein region importance scores to obtain layer-wise residue importance scores,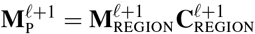, and ligand functional group importance scores to obtain layer-wise atom importance scores,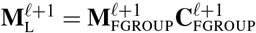.

#### Interpretable Fingerprint

During *ℒ* iterations of PGC layers, PSICHIC accumulates the residue and atom importance scores from each layer. After that PSICHIC learns strictly positive weight vectors to project the layer-wise scores into the final importance score for residues,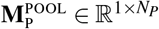, and atoms,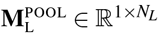:

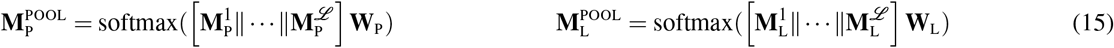

where ∥represents concatenation operation on the layer-wise importance scores, and **W**_P_ and **W**_L_ are the parameters of linear projections for protein and ligand, respectively. Importantly, PSICHIC constrains the parameters of the linear projections to be positive by initializing them as log-parameterized weights. Consequently, a higher importance in PGC layers will only lead to a higher final importance score,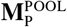 and 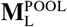. This ensures that the final importance scores of the ligand atoms and protein residues are directly determined by the magnitude of intermolecular interactions within the network.

Using the importance scores, PSICHIC pools the protein residues into a single protein feature weighted by their relative importance scores,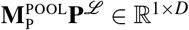, and ligand atoms into a single ligand feature weighted by their relative importance scores,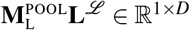. Finally, PSICHIC obtains the interaction fingerprint, **f**, by concatenating the protein and ligand feature after projecting them into a joint interaction space using MLPs:

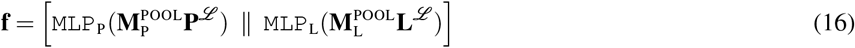

#### Application Layer and Objective Functions

Under the standardized backbone, PSICHIC is trained using application-specific MLP(s) on the interaction fingerprint, **f**. We optimize PSICHIC with supervised objective(s), **O**_APPLICATION(S)_, based on the type of interaction properties PSICHIC aims to predict. For binding affinity value prediction on PDBBind datasets, we have BINDINGAFFINITY = MLP_affinity_(**f**) and PSICHIC is optimized using mean-squared error loss (MSE), **O**_APPLICATION(S)_ = **O**_MSE_. For binary activity classification and three-way functional effect classification, we use CLASS = MLP_classification_(**f**). In this case PSICHIC is optimized using cross-entropy loss, **O**_APPLICATION(S)_ = **O**_CE_. This is applied to the binary active classification datasets (Human, BioSNAP, and BindingDB) as well as the protein-ligand functional effect dataset that we curated. For large-scale datasets, PSICHIC is jointly trained to predict both binding affinity and three-way functional effect classes. Consequently, PSICHIC has two MLP heads for binding affinity and functional effect class, respectively. Both heads share the same fingerprint, and the supervised objective is: **O**_APPLICATION(S)_ = **O**_MSE_ + **O**_CE_. Finally, as PSICHIC also learns to cluster protein region at every PGC layer during training, PSICHIC is also optimized with the unsupervised MinCUT loss, **O**_MINCUT_52, together with the supervised objectives. Since every layer in the PGC Layer has a clustering loss, the total MinCut loss is computed as 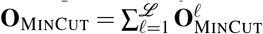. Together, the optimization objective of PSICHIC is: **O**_PSICHIC_ = **O**_APPLICATION(S)_ + **O**_MINCUT_.

### PSICHIC Optimization

#### Training PSICHIC

To train PSICHIC, we used Adam^56^ with *β*_1_ = 0.9, *β*_2_ = 0.999, and *ε* = 10^*−*8^ at a static learning rate of 10^*−*4^. We clipped the global norm of the gradient at 1.0 and applied a weight decay of 10^*−*4^ to provide a small amount of regularization^57^. In all experimental settings, PSICHIC used the same optimizer configurations, except for PDBBind v2016 and the large-scale interaction dataset due to the size of the training data. The PDBBind v2016 set has only 3,390 training samples, so we modified *β*_2_ = 0.99, and *ε* = 10^*−*5^ to limit the parameter-wise learning rate. We set the learning rate to 10^*−*5^ for the large-scale interaction dataset due to the small batch size and data distribution imbalance, which could lead to learning instability (Extended Data Fig. 8). All experiments used a virtual machine with an NVIDIA A100 Tensor Core GPU. **Pretrain-then-finetune Multi-task PSICHIC**. To train multi-task PSICHIC on large scale interaction dataset, we implemented a two-stage, pretrain-then-finetune strategy. In the pre-training stage, we categorized protein-ligand interactions into groups such as ProteinA_Binder and ProteinA_Non-Binder during the pre-training stage. Sampling weights were assigned to each group based on the square root of its sample size and capped at the 90th percentile to prevent overrepresentation. Despite these measures, the agonist and antagonist classes were underrepresented (see Extended Data Fig. 8). To address this, the second fine-tuning stage exclusively trains the functional effect classification head of PSICHIC on agonist and antagonist interactions involving A_1_R and proteins with over 50% sequence overlap, as defined by MMSeq^58^. This approach aimed to optimize PSICHIC for targeted A_1_R virtual screening without sacrificing knowledge gained from the diverse pre-training data. **Hyperparameters of PSICHIC**. Hyperparameters were tuned using PDBBind v2020 benchmark dataset (see Supplementary Methods 1). Consequently, we fix the number of layers, *ℒ*, to be 3, and the number of protein clusters, *K*, to be {5, 10, 20 }for the three layers respectively. We set *K* = 20 for the final layer because the ratio of total protein residues to pocket residues is approximately 20^59^. We find it necessary to gradually increase *K* by setting the first two layers to be *K* = 5 and 10 for MinCUT objective^52^, **O**_MINCUT_, to converge. The hidden dimension, *D*, of PSICHIC is set to be 200. All MLPs have a single hidden layer with dimensions set to be 2 ×*D*, and an output layer to be *D*. We use ReLU activation function^60^. Lastly, we utilize 5 multi-head attention mechanisms^61^ and apply a 0.2 dropout rate on the attention logits before softmax activations of the intermolecular forces (Eq. 9 and Eq. 10) in PGC layers to prevent the inferred protein-ligand interactions from getting stuck in local optima. To prevent over-tuning hyperparameters, the same configuration is then applied across all experimental settings.

### Pharmacological Materials

Chinese hamster ovary (CHO) Flp-IN™ (FlpInCHO) cells, Dulbecco’s modified Eagle’s medium (DMEM) were purchased from Invitrogen (Carlsbad, CA). DMEM high glucose medium was purchased from Sigma-Aldrich. Fetal bovine serum (FBS) was purchased from ThermoTrace (Melbourne, VIC, Australia). Hygromycin B was purchased from Thermofisher. Adenosine deaminase (ADA) was purchased from Roche (Basel, Switzerland). LANCE®Ultra cAMP kit was purchased from PerkinElmer (Boston, MA). 5’-(N-Ethylcarboxamido)adenosine (NECA), Forskolin, Rolipram, and Dimethyl sulfoxide (DMSO) were purchased from Sigma-Aldrich. Compounds within the MIPS library were purchased from commercial vendors (Chembridge, Lifechemicals, Mol Port).

### Cell culture

FlpInCHO cells stably transfected with human A_1_R (A_1_R-FlpInCHO) were generated as previously describe^62^ and maintained in DMEM supplemented with 10% FBS and 500 *µg/mL* hygromycin-B. Non-transfected FlpInCHO cells (NT-FlpINCHO) were maintained in DMEM supplemented with 10% FBS. Both cell lines were grown at 37°C in a humidified incubator containing 5% CO2. Cells were expanded and froze down in 1 mL aliquots (10 x 106 cells/mL) in freezing media (FBS, 10% DMSO) and stored at -80°C.

### cAMP accumulation assay

A_1_Rs preferentially couple to Gi/o proteins to decrease 3’5’-cyclic adenosine monophosphate (cAMP) accumulation. A_1_R-FlpINCHO cells are stimulated with forskolin to increase intracellular cAMP. The LANCE®Ultra cAMP kit was used to measure the cAMP produced by the A_1_R cells. TR-FRET signal is detected using a PHERAstar FSX (BMG LABTECH) plate reader. The screening process to identify A_1_R agonists can be split into three parts: primary high throughput screening, confirmation and counter-screening. The high-throughput screening was carried out at the National Drug Discovery Centre, WEHI, in Parkville, Australia, with support from the Australian Government Medical Research Future Fund (MRFF); confirmed hits were validated at MIPS.

In the primary high-throughput screening campaign, the objective was to identify A_1_R agonists using A_1_R-FlpINCHO cells. The screen utilized the MIPS library, which contains 30,282 compounds. To assess the decrease in forskolin-induced cAMP accumulation, we employed the LANCE® Ultra cAMP assay in conjunction with an EC25 concentration of the A_1_R orthosteric agonist NECA. The assay was performed in a 1536-well microplate format the total reaction volume was 6 µL per well (2 µL cells + 2 µL Forskolin + 2 µL LANCE cAMP detection reagents). For detection reagent mixture, the ratio for Eu-cAMP and Ulight anti-cAMP were 1:500 and 1:1500, respectively. These were then transferred to an ultra-high-throughput biochemical screening platform. In this primary screen, the compounds were tested at a final concentration of 12.5 µM. The assay achieved an average signal-to-background ratio of approximately 3.5 and a Z’ factor of around 0.6. Hits were selected based on the criterion of compounds exhibiting a percentage inhibition greater than three standard deviations above the mean of the samples in each batch.

For confirmation, counter-screening, and final hit list compilation, we conducted a 10-point dose-response test using 1536-well microplate format. These tests were performed using both A_1_R-FlpINCHO and NT-FlpINCHO cells (the counter-screen assay was used to identify compounds that generated false-positive results in the primary assay). Compounds were titrated starting from a concentration of 100 µM, using a 1:2 dilution series. In the primary assay, we achieved an average signal-to-background ratio of 6.2; in the counter-screen assay, this ratio was 8.8. The corresponding average Z’ values for the primary assay and the counter-screen assay were 0.57 and 0.7, respectively. Ultimately, three compounds (MIPS1, MIPS2, and MIPS3) were identified as being active in the primary assay and as being inactive in the counter-screen assay. Agonist activity was evaluated for three hit compounds at a final concentration of 10 µM using a 384-well format. Tests were conducted on both A_1_R-FlpINCHO and non-transfected FlpINCHO cells. MIPS1 induced an inhibition of cAMP accumulation that was comparable to the effect of the agonist, NECA, and showed no effect on non-transfected cells.

## Data Availability

All raw and benchmark data resources used are publicly available. They were obtained from the following databases: Protein Data Bank (https://www.rcsb.org), UniProt (https://www.uniprot.org), PDBBind (http://www.pdbbind.org.cn/), ExCAPE-ML (https://solr.ideaconsult.net/search/excape/), and Papyrus (https://zenodo.org/record/7019874). While Cortellis Drug Discovery is not publicly available, the manually curated protein-ligand functional effect dataset and large-scale interaction dataset used for training and testing the models are fully available from the GitHub repository referenced below.

## Code Availability

A GitHub repository containing the source code and data files for retraining and evaluating PSICHIC is available at https://github.com/huankoh/PSICHIC. The repository contains a user-friendly, open-source online platform for PSICHIC’s virtual screening application, integrated with Google Colaboratory for easy web-based interaction. The weights of the trained PSICHIC model are also available in the repository.

## Acknowledgements

Research on adenosine receptor signalling was supported by a National Heart Foundation Future Leader Fellowship (101857 [L.T.M]), National Health and Medical Research Council (NHMRC) of Australia Ideas grant (APP2013629 [L.T.M, G.W, A.T.N.N]), and a Department of Health and Aged Care (MRFF) Stem Cell Therapies Mission grant (MRF2015957, [L.T.M, A.T.N.N]). H.Y.K scholarship is supported by the Australian Government Research Training Program (RTP) Scholarship and the Australian Research Council under grant ARC DP210100072. High throughput screening was performed at the National Drug Discovery Centre, WEHI, Parkville, Australia, with support from the Australian Government Medical Research Future Fund (MRFF). Our acknowledgment extends to Cortellis Drug Discovery Intelligence for granting public access to the curated functional effect dataset. Special thanks to Monash Institute of Pharmaceutical Sciences (MIPS) Monash University for access to the MIPS library, in particular Patrick Sexton and Arthur Christopoulos for purchase of the MIPS library and to Jonathan Baell for the design of the library. We wish to thank Cam Sinh Lu for assistance with pharmacological evaluation. Computational resources were generously provided by the Nectar Research Cloud, a collaborative Australian research platform supported by the NCRIS-funded Australian Research Data Commons (ARDC) and the MASSIVE HPC facility.

## Author contributions

H.Y.K. designed and developed the PSICHIC method, evaluated PSICHIC against leading methods, and applied it to virtually screen the MIPS library for novel A_1_R agonists. A.T.N.N performed the pharmacological validation of novel A_1_R agonists and conducted data analysis. H.Y.K. and A.T.N. prepared figures and wrote the manuscript. A.T.N.N, S.P., L.T.M., and G.W. supervised the project. Project design, data interpretation, and manuscript preparation were performed by all authors.

## Competing interests

The authors declare no competing interests.

**Extended Data Fig. 1.**
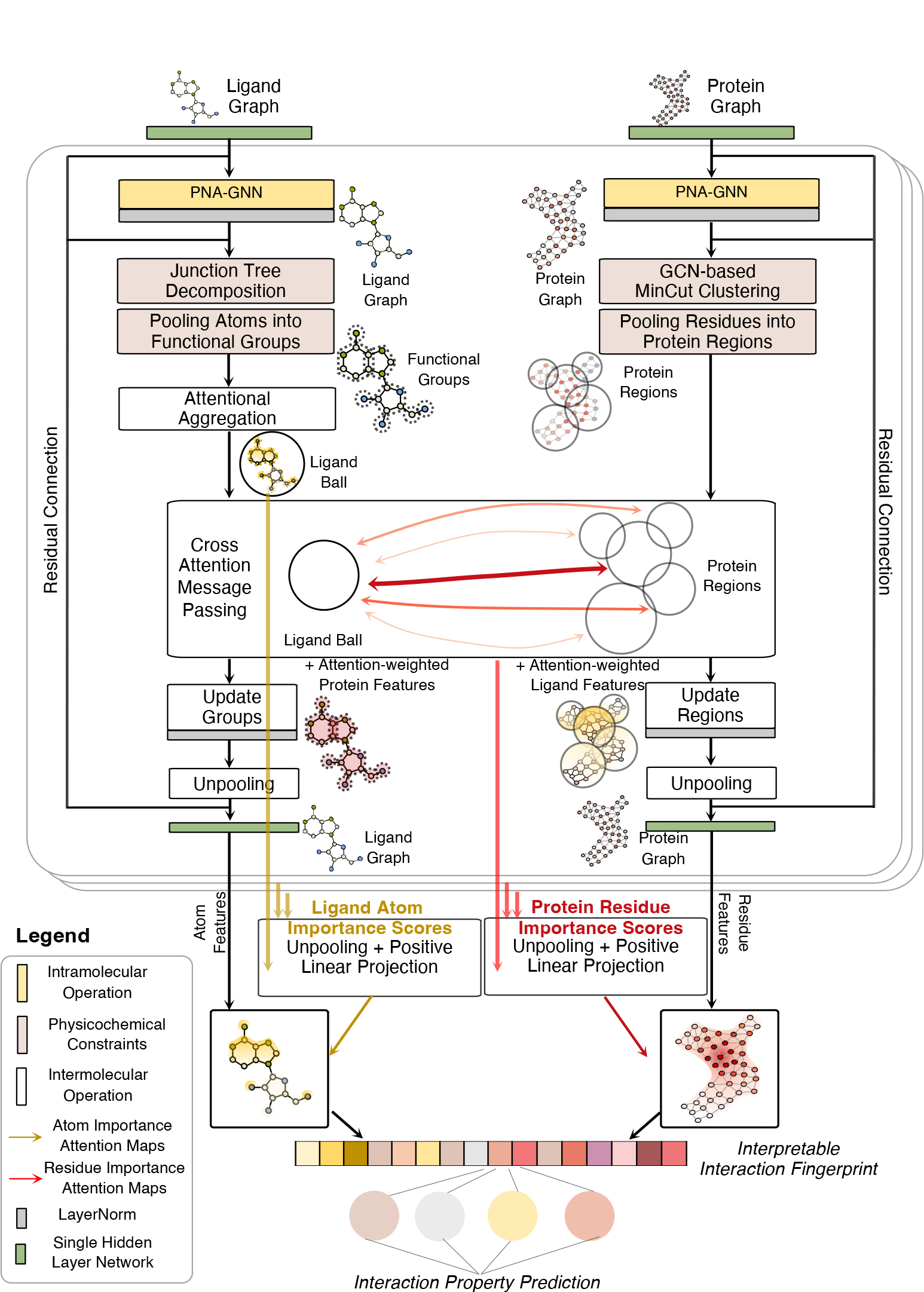
Schematic Diagram of PSICHIC Architecture. The figure should be read from top to bottom. Initially, ligand atom and protein residue graphs pass through a single hidden layer network before entering three physicochemical graph convolutional layers. Within each layer, intramolecular forces (represented by yellow blocks) are modeled using two independent PNA-GNNs^26^. PSICHIC then imposes physicochemical constraints (light red blocks) by pooling ligand atoms and protein residues using junction tree decomposition^54^ and MinCut Clustering^52^, respectively. PSICHIC models intermolecular forces (white blocks) in three steps: first, it aggregates ligand functional groups into a ‘ligand ball’ using attentional aggregation; second, it models the interaction strengths between this ligand ball and protein regions through cross-attention message passing, where features are weighted and transferred between the ligand and protein; third, PSICHIC unpools the functional groups and clustered regions back into updated ligand atoms and protein residues. Finally, PSICHIC generates an interaction fingerprint that weights atoms and residues based on importance scores from intermolecular forces. This fingerprint feeds into a single-hidden-layer network for predicting interaction properties. The figure can be read in conjunction with the Method section.

**Extended Data Fig. 2.**
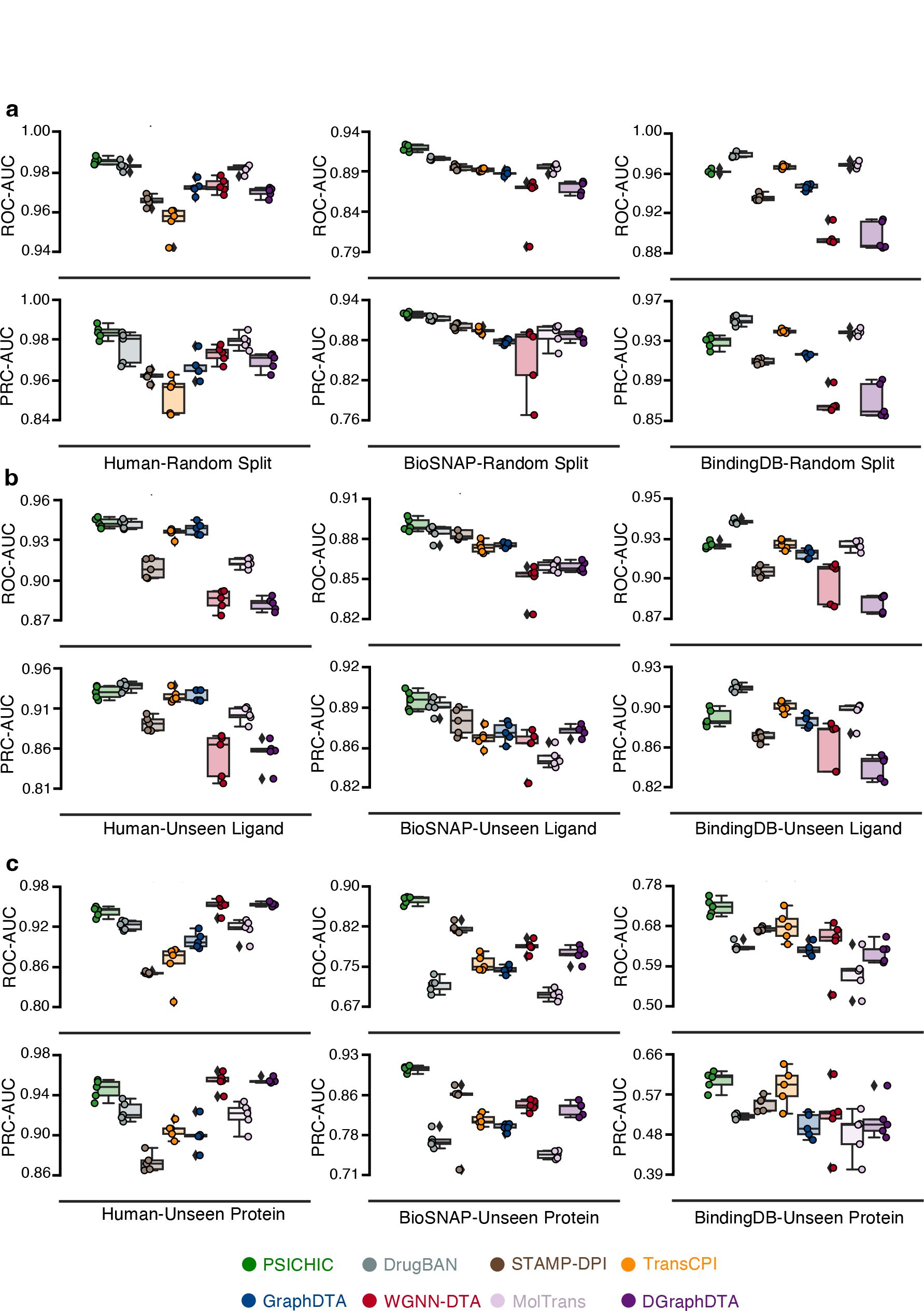
Performance of Sequence-Based Methods on Human, BioSNAP, and BindingDB Datasets. Using sequence-only datasets from Human^36^, BioSNAP^37^, and BindingDB^38^ as benchmarks, we applied three splitting strategies to evaluate sequence-based methods: (**a**) random split, (**b**) unseen ligand split, and (**c**) unseen protein split. The unseen ligand split ensured that ligand scaffolds in the test sets were unseen during model training, while the unseen protein split ensured that protein targets in the test sets were likewise unseen during training. AUC-ROC and AUC-PRC stand for the Area Under the Receiver Operating Characteristics Curve and Area Under the Precision-Recall Curve, respectively. Models are arranged in the boxplots according to their average AUC-ROC performance rankings for sequence-based methods (see Supplementary Table 4 for robust pairwise model comparisons through the Multiple Comparison Matrix^63^).

**Extended Data Fig. 3.**
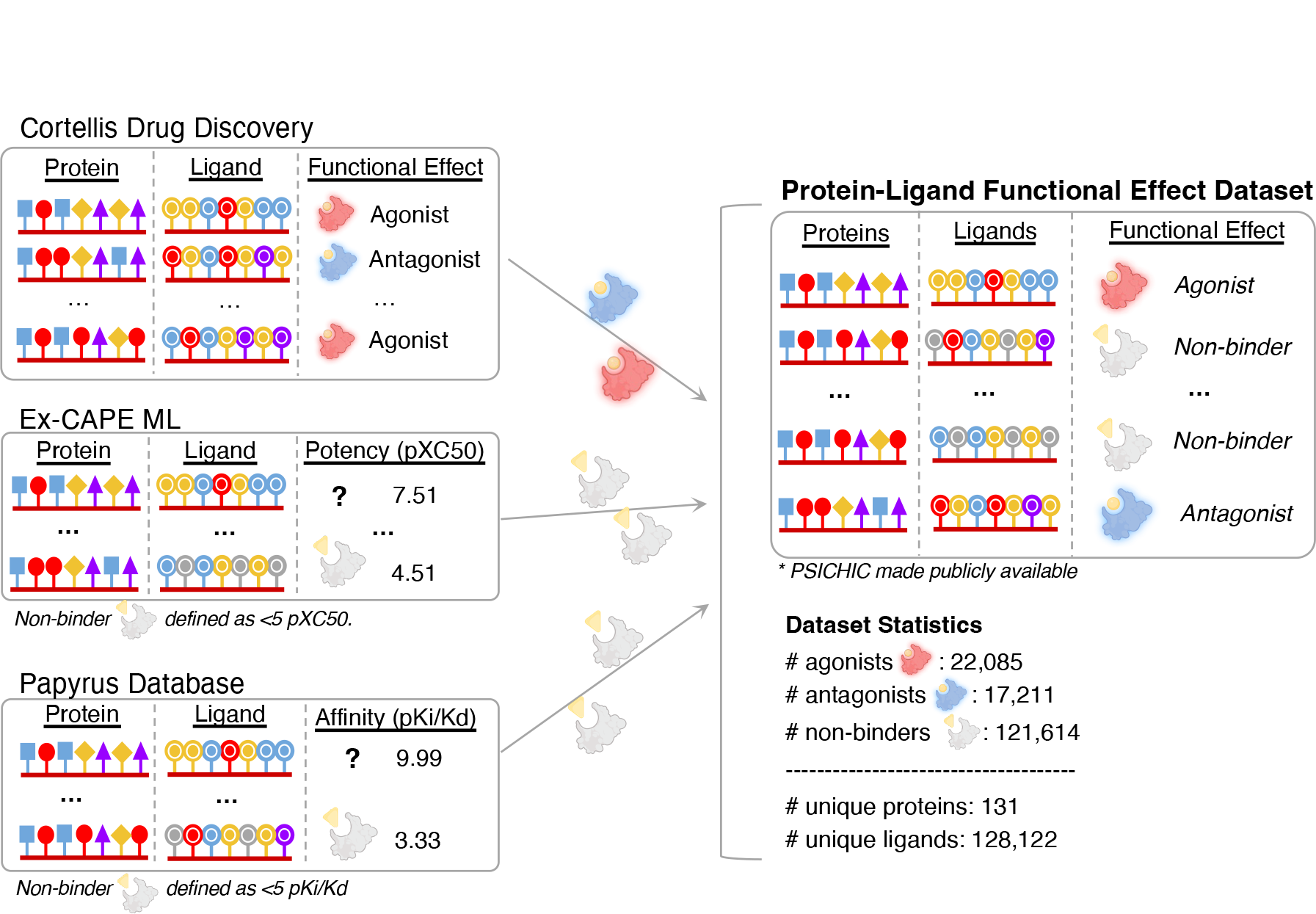
High-Quality Protein-Ligand Functional Effect Dataset Curation. As a data-driven framework, PSICHIC can predict various interaction properties based on labeled sequence datasets. To enable PSICHIC in predicting ligand-protein functional effects, we curated data from reputable databases, specifically Cortellis Drug Discovery^64^, ExCAPE-ML^41^, and Papyrus^42^. We gathered samples from Cortellis on 02/02/2023, focusing on the proteins categorized as ‘Receptor’ in the database as well as protein receptors with over 20 samples. This resulted in 22,085 agonists and 17,211 antagonists [Source: Cortellis Drug Discovery Intelligence, 02 02, 2023 https://www.cortellis.com/drugdiscovery/ ©2023 Clarivate. All rights reserved.]. Due to the absence of negative data, we adopted a rigorous approach to include decoys. From ExCAPE-ML^41^ and Papyrus^42^, we selected protein-ligand pairs with pXC50 or pKi/pKd values below 5, focusing on high-quality data in the database. After standardizing the molecular data using the ChEMBL pipeline^65^, we generated a dataset comprising 160,910 unique protein-ligand pairs: 22,085 agonists, 17,211 antagonists, and 121,614 non-binders, which includes 131 unique protein receptors and 128,122 unique ligands. The dataset also contains assay-dependent potency values (EC50 for agonists and IC50 for antagonists), which were omitted for model training. Instead, they were used in the UMAP plot of Fig. 2. Along with this study, the curated dataset is publicly available.

**Extended Data Fig. 4.**
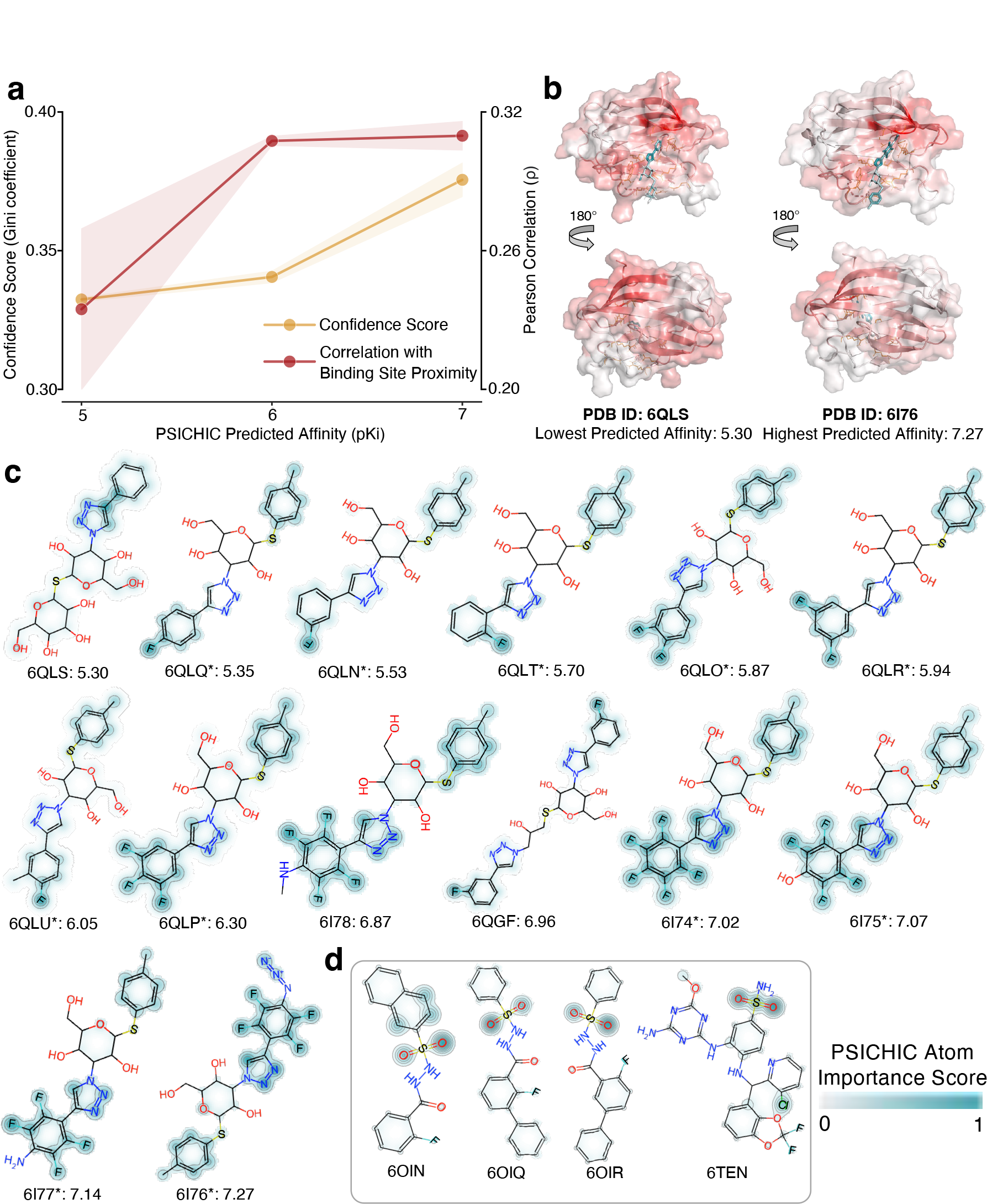
Case Study of Pharmacophore Analysis on Galectin-3 Target Protein. **a**, The line plot illustrates relationships based on the predicted binding affinity by PSICHIC: (1) an orange line shows the relationship between affinity (x-axis) and confidence in residue importance scores (y-axis), the latter being measured by the Gini coefficient. (2) a red line depicts the relationship between affinity (x-axis) and the correlation of PSICHIC residue importance scores with residues’ proximity to the binding site (y-axis). The error band (shaded region) of line plots indicates one standard error of the mean (SEM). **b**, Complex structures of Galectin-3 with low (PDB ID: 6QLS) and high (PDB ID: 6I76) predicted binding affinities are displayed. A darker red hue on the structures signifies higher PSICHIC importance scores. For low-affinity 6QLS, scores are dispersed—sometimes extending 180 degrees away from the binding site—yet key binding residues are highlighted. Conversely, scores for high-affinity 6I76 are primarily focused on the binding sites. **c**, PSICHIC’s ligand atom importance scores spotlight key functional groups, especially fluorine atoms, in their interaction with Galectin-3. The 14 binding ligands, 11 of which share the same scaffold (*), are ordered based on PSICHIC-predicted affinities. The emphasis of PSICHIC on the fluorine functional groups aligns with the original studies, where fluorine plays a critical role in forming fluorine–amide interactions with Galectin-3’s binding site^66, 67^. **d**, As a control, PSICHIC did not universally prioritize fluorine atoms when the fluorine functional groups do not form important interactions with the target protein. Further details are provided in Supplementary Methods 8.

**Extended Data Fig. 5.**
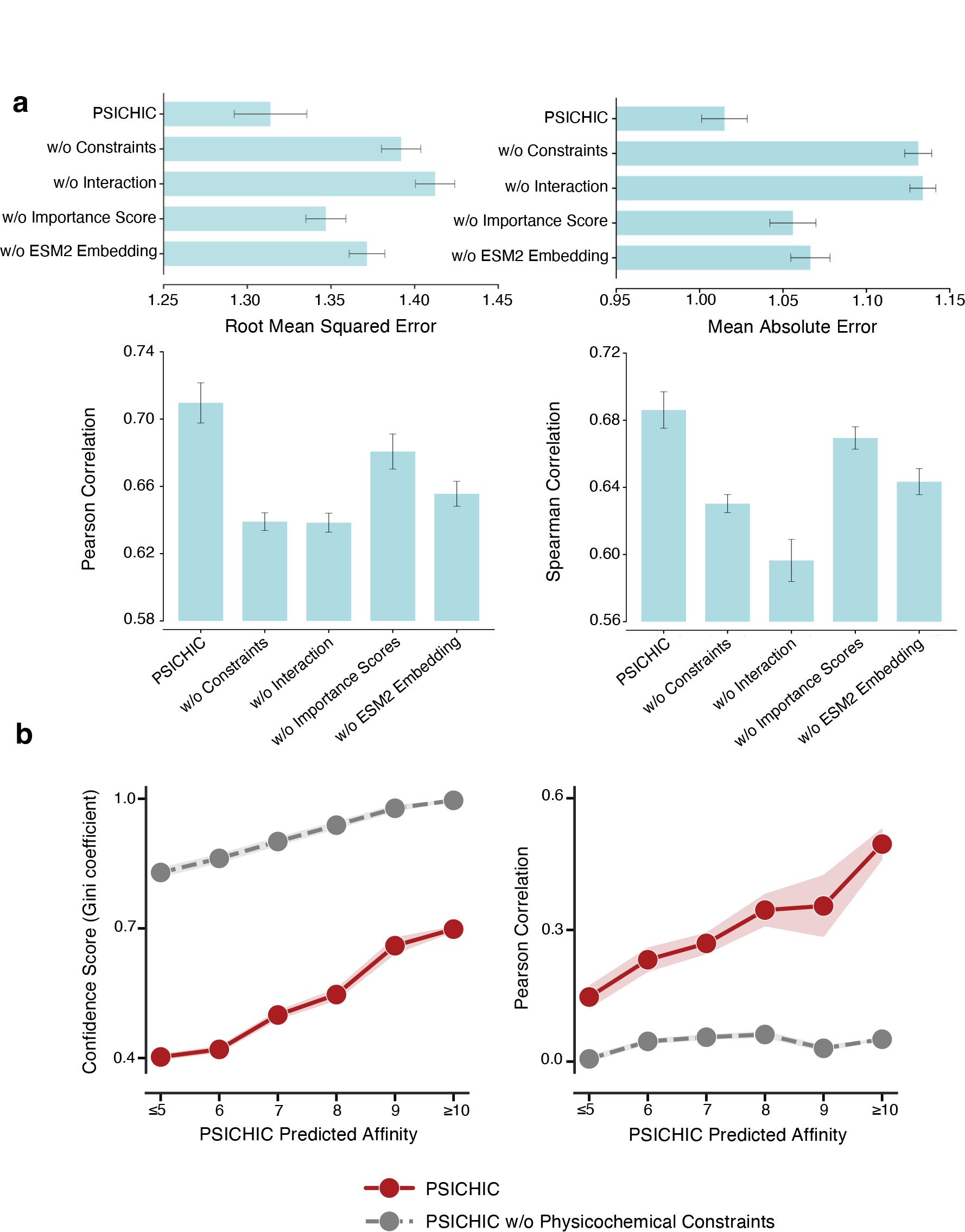
Ablation study of PSICHIC’s Architecture on PDBBind v2020 benchmark dataset. Removing PSICHIC’s physicochemical constraints (w/o Constraints) resulted in diminished performance. Omitting explicit interactions (w/o Interaction) involved modeling only intramolecular forces and implicitly capturing interactions by concatenating both protein and ligand representations into a fingerprint. Interestingly, removing physicochemical constraints yielded results almost identical to those obtained when not modeling interactions at all, highlighting the importance of incorporating constraints. Removing the Importance Score mechanism (w/o Importance Score) made the model less accurate. Finally, the absence of ESM2 embedding led to slight underperformance compared to TankBind^8^, which uses its own evolutionary residue embeddings from the TAPE protein model^68^. This comparison is not directly comparable but favors TankBind. **b**, Physicochemical constraints were central to PSICHIC’s ability to learn patterns of protein-ligand interactions that adhere to physicochemical principles. The left plot shows that without constraints, PSICHIC exhibits *stronger* confidence in residue importance scores. The right plot indicates that this confidence is misplaced because the scores do not correlate with the proximity of the residues to the binding site. This suggests that sequence-based methods could overfit the data if physicochemical constraints are not incorporated.

**Extended Data Fig. 6.**
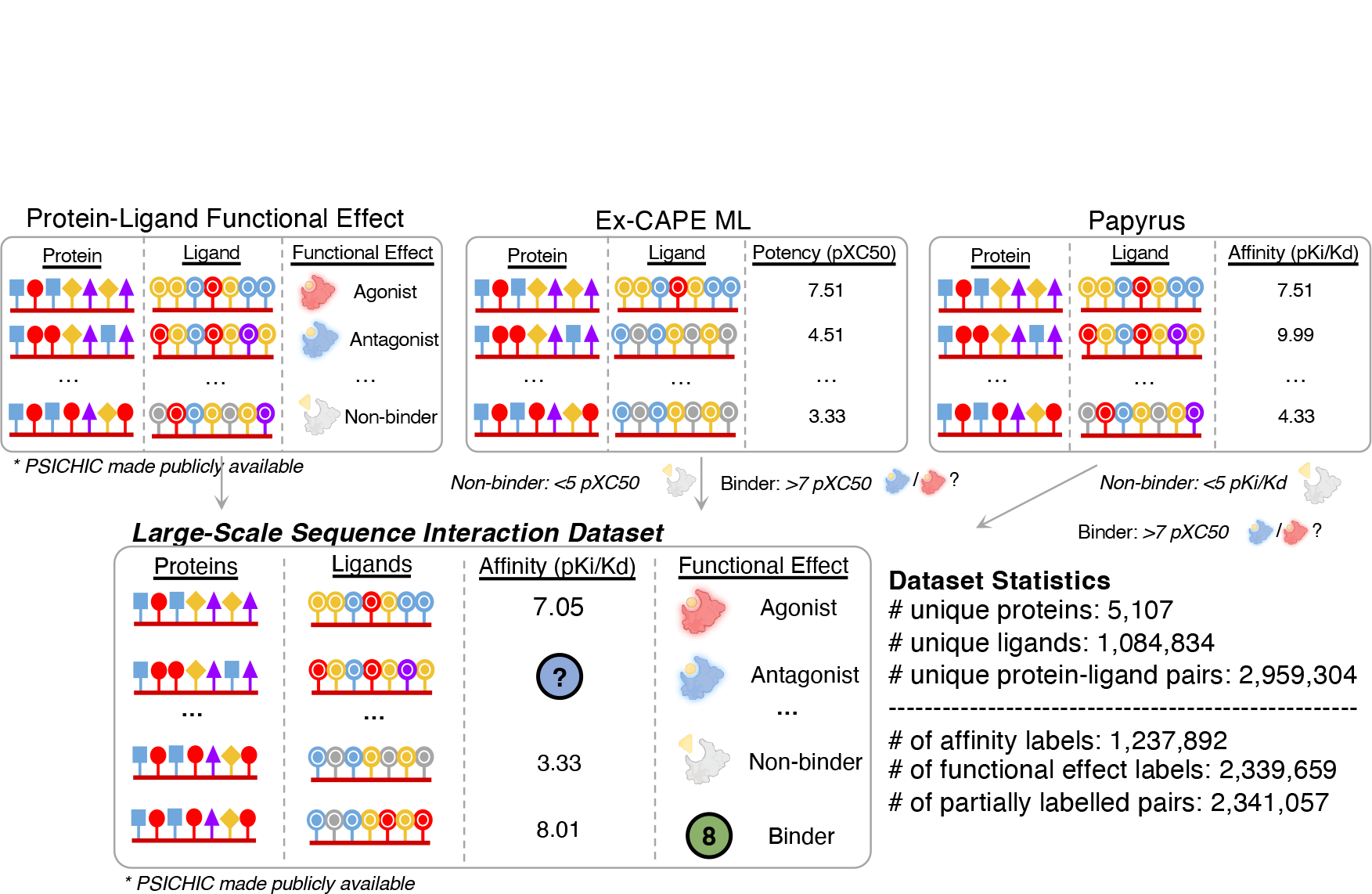
Large Scale Interaction Dataset. From the Protein-Ligand Functional Effect Dataset, we extracted samples comprising a protein, ligand, and functional label (agonist, antagonist, or non-binder; Extended Data Fig. 3, Supplementary Methods 9). From ExCAPE-ML^41^, protein-ligand pairs with pXC50 values below 5 were labeled as non-binders and above 7 as binders. From Papyrus^42^, we selected pairs with high-quality binding affinity, labeling those with pKi/Kd below 5 as non-binders and above 7 as binders. These databases were standardized and normalized using the ChEMBL pipeline^65^ and combined to form a large-scale dataset of approximately 3 million unique protein-ligand pairs, labeled with either binding affinity or functional effect. The final dataset contains 618,247 fully labeled and 2,341,057 partially labeled pairs, encompassing 5,107 unique proteins and 1,084,834 unique ligands.

**Extended Data Fig. 7.**
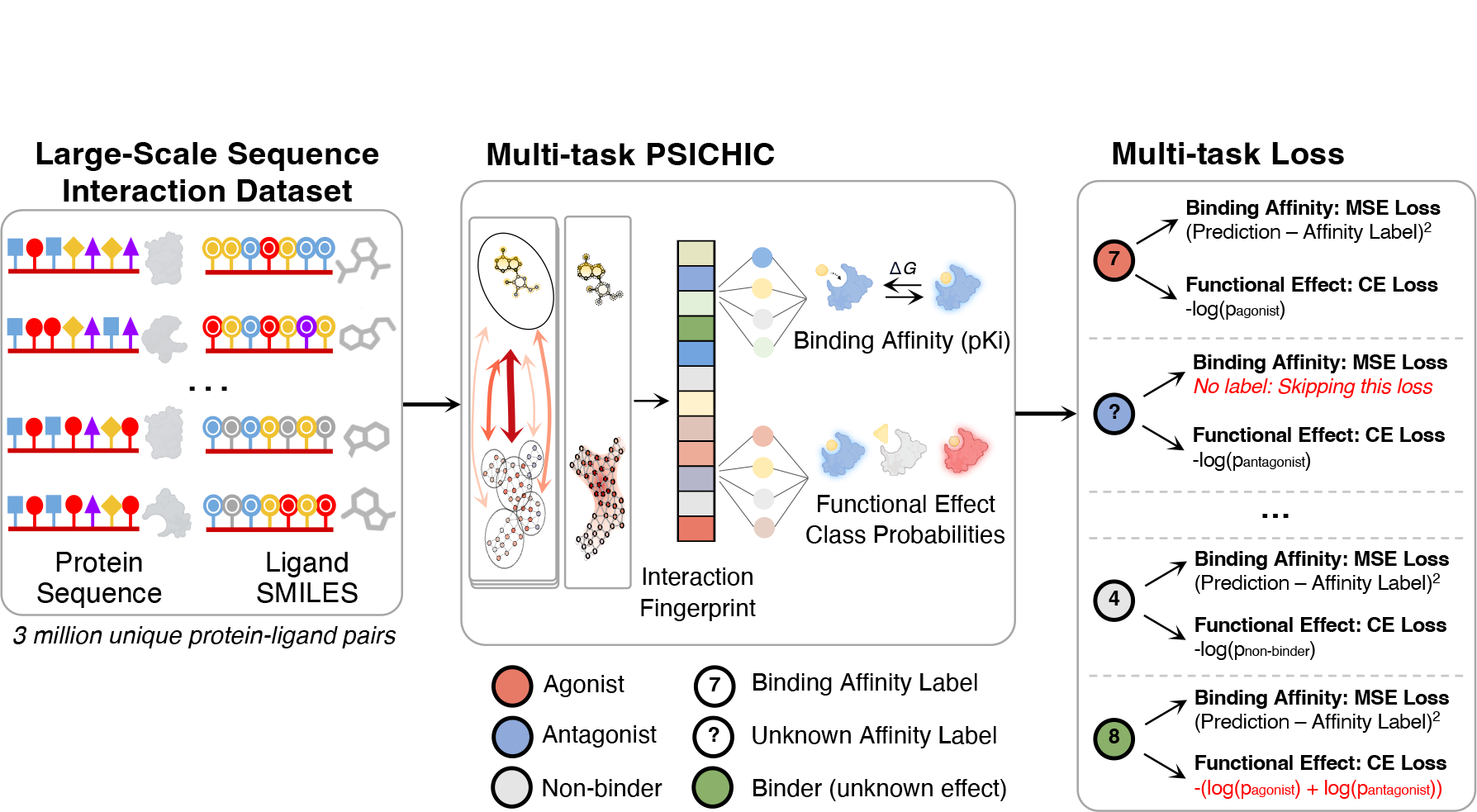
Multi-task optimization of PSICHIC. Schematic illustrating the multi-objective loss function for handling partially labeled protein-ligand pairs in the large-scale interaction dataset we curated. Two scenarios of partial annotation were addressed: (i) when only the functional effect class (agonist, antagonist, or non-binder) is known, only the cross-entropy loss is calculated; (ii) when only binding affinity is known and ligand is a binder (i.e., functional effect of the ligand on its protein target is not known), the cross-entropy loss function is minimized as −log(*p* ≠ non-binder) or − (log(*p*_agonist_) + log(*p*_antagonist_)), together with the mean-squared-error loss. Refer to Methods and Supplementary Methods 9 for full details on training PSICHIC on large-scale partially annotated dataset.

**Extended Data Fig. 8.**
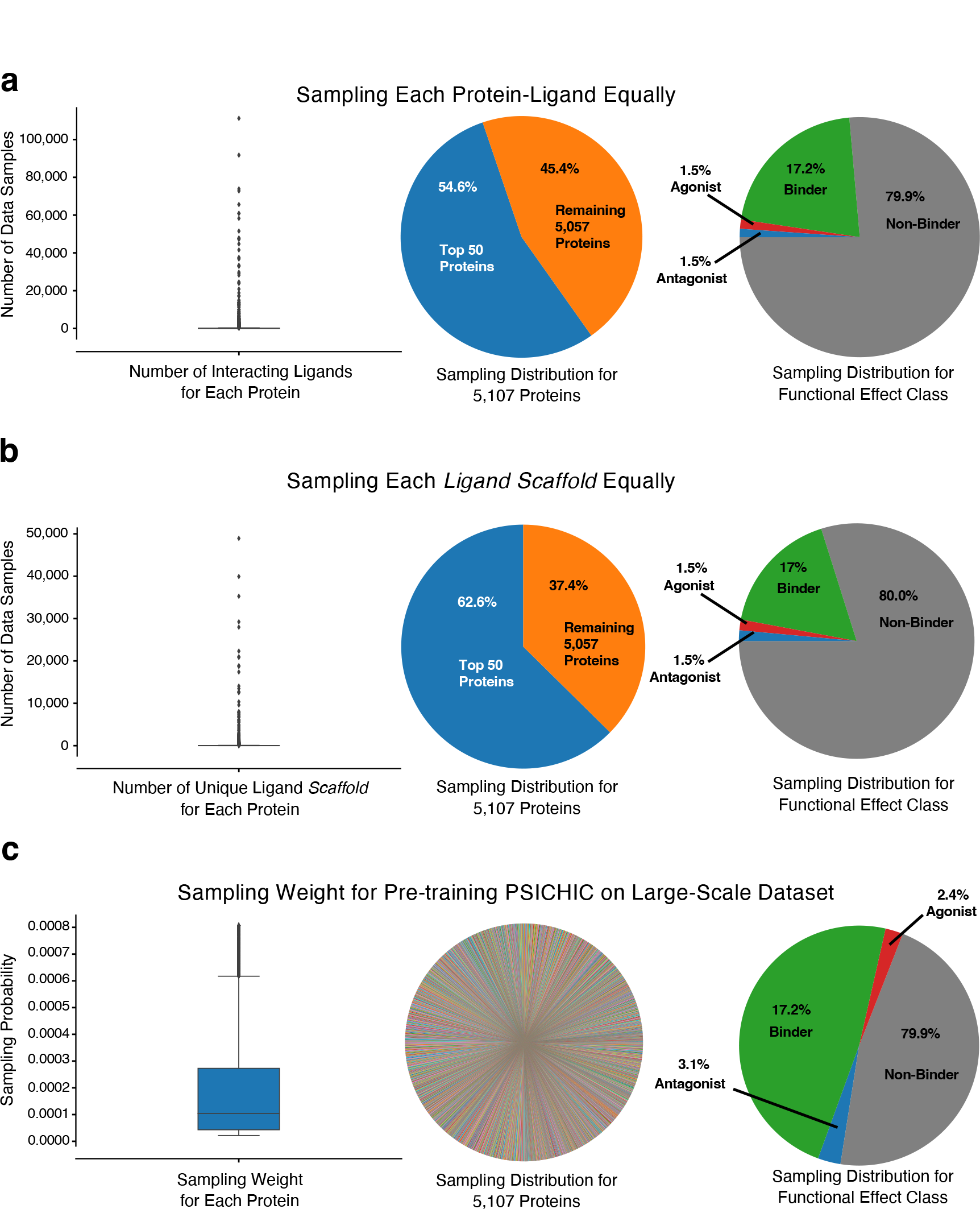
Diverse protein-ligand interaction sampling strategy. **a**, The box plot and pie charts demonstrate that, despite the dataset’s extensive scale and diversity, it was notably imbalanced. Specifically, the top 50 proteins (1% of 5,107) accounted for over half of all samples, most of which were non-binders. This imbalance risked overfitting when training PSICHIC through a random sampling strategy. **b**, The issue persisted even when each ligand scaffold was sampled equally, as illustrated by the box plot and pie charts. **c**, To mitigate this, we implemented a two-stage, pretrain-then-finetune strategy for PSICHIC. During pre-training, we first categorized each protein-ligand interaction sample based on its protein and whether it was a binder or a non-binder. Specifically, we created groups like ProteinA_Binder, ProteinA_Non-Binder, …, ProteinZ_Binder, ProteinZ_Non-Binder. We then assigned a sampling weight to each group proportional to the square root of the group’s size, capping this at the 90th percentile to prevent overrepresentation. Within each group, we ensured that each ligand scaffold was sampled equally. The resulting sampling distribution for the pre-training of PSICHIC is illustrated in the box plot and pie charts. Nonetheless, the agonist and antagonist classes remained underrepresented during pre-training. Hence, during the fine-tuning stage, we trained only the functional effect classification head of PSICHIC exclusively on agonist and antagonist interactions involving A_1_R and related proteins. Related proteins are defined as proteins greater than 50% sequence overlap, according to MMSeq^58^. This approach ensured that PSICHIC was optimized for a targeted virtual screening, while preserving the richness of knowledge gained from the diverse training data. See Supplementary Methods 9 for full details.

**Extended Data Table 1.** Table Summary describing the size, split and interaction property prediction task of the datasets in the benchmark.

| **Benchmark Dataset** | **Interaction Property Type** | **Train/Valid/Test Sample Count** | **Description** |
| --- | --- | --- | --- |
| PDBBind v2016 | Binding Affinity | 3,390 / 377 / 290 | Standard split with test set contains high-quality complexes from PDBBind Core Set<sup>6,28</sup>. |
| PDBBind v2020 | Binding Affinity | 17,797 / 968 / 363 | Standard split using temporal split (Training data include PDB entries before 2019, and entries during and after that point are used as test data)<sup>8</sup>. |
| Human: Random Split | Binary Presence of Interaction | 4,197 / 600 / 1,200 | Samples were randomly split. |
| Human: Unseen Ligand Scaffold Split | Binary Presence of Interaction | 4,197 / 599 / 1,201 | Ligand scaffolds were split to ensure that the scaffolds in the test (and validation) set were not seen during training. |
| Human: Unseen Protein Split | Binary Presence of Interaction | 4,022 / 600 / 1,375 | Protein targets were split to ensure that the targets in the test (and validation) set were unseen during training. |
| BioSNAP: Random Split | Binary Presence of Interaction | 19,224 / 2,747 / 5,493 | Samples were randomly split. |
| BioSNAP: Unseen Ligand Scaffold Split | Binary Presence of Interaction | 19,224 / 2,746 / 5,494 | Ligand scaffolds were split to ensure that the scaffolds in the test (and validation) set were unseen during training. |
| BioSNAP: Unseen Protein Split | Binary Presence of Interaction | 18,841 / 2,565 / 6,058 | Protein targets were split to ensure that the targets in the test (and validation) set were not seen during training. |
| BioSNAP: Unseen Protein and Ligand Split | Binary Presence of Interaction | 9,766 / 3,628 / 907 | Similar to DrugBAN, we split the dataset so that both proteins and ligands in the test set are unseen and substantially different from those in the training set. Single-linkage clustering with a minimum distance threshold of 0.5 is used to separate ligand compounds (ECFP4 feature) and target proteins (integral PSC feature). |
| BindingDB: Random Split | Binary Presence of Interaction | 34,439 / 4,920 / 9,840 | Samples were randomly split. |
| BindingDB: Unseen Ligand Split | Binary Presence of Interaction | 34,439 / 4,918 / 9,842 | Ligand scaffolds were split to ensure that the scaffolds in the test (and validation) set were not seen during training. |
| BindingDB: Unseen Protein Split | Binary Presence of Interaction | 34,236 / 3,909 / 11,054 | Protein targets were split to ensure that the targets in the test (and validation) set were not seen during training. |
| BindingDB: Unseen Protein and Ligand Split | Binary Presence of Interaction | 14,928 / 7,114 / 1,779 | Similar to DrugBAN, we split the dataset so that both proteins and ligands in the test set are unseen and substantially different from those in the training set. Single-linkage clustering with a minimum distance threshold of 0.5 is used to separate ligand compounds (ECFP4 feature) and target proteins (integral PSC feature). |
| PSICHIC: Functional Effect Dataset* | Protein-Ligand Functional Effects | 112,637 / 16,091 / 32,182 | Samples were randomly split. |
| PSICHIC: Large-scale Interaction Dataset*+ | Binding Affinity and Functional Effects | 2,959,304 / 0 / 0 | Samples were used for training multi-task PSICHIC. |

\* Curated by us, and will be made publicly available.
+ Large-scale interaction dataset is used to train PSICHIC for real-world applications.

**Extended Data Table 2.** Performance comparison on the PDBBind v2016 and v2020 datasets. Prediction errors are measured using Root Mean Square Error (RMSE) and Mean Absolute Error (MAE). Correlation with experimental affinity values is determined using Pearson and Spearman correlation coefficients. The r_m_^2^, which quantifies the closeness between observed and predicted activity^69^, and the Concordance Index (CI), which assesses the model’s ability to rank protein-ligand pairs, are also reported. An upward arrow (↓) denotes higher scores are better, and a downward arrow (↑) denotes the reverse. Average results and standard deviations (in parentheses) from five independent runs are reported. The best performance for each metric is highlighted in **bold**, while the second-best performance is underlined.

|  | Methods | Hi-res<br>Structure | Co-crystal<br>Complex | PDBBind v2016 |  |  |  |  |  |
| --- | --- | --- | --- | --- | --- | --- | --- | --- | --- |
| | | | | RMSE ↓ | MAE ↓ | Pearson ↑ | Spearman ↑ | $r_m^2$ ↑ | CI ↑ |
| Complex | Pafnucy<sup>30</sup> | ✓ | ✓ | 1.565 (0.023) | 1.261 (0.017) | 0.699 (0.008) | 0.693 (0.007) | 0.458 (0.03) | 0.751 (0.004) |
|  | OnionNet<sup>31</sup> | ✓ | ✓ | 1.377 (0.040) | 1.092 (0.029) | 0.789 (0.019) | 0.789 (0.017) | 0.558 (0.035) | 0.798 (0.008) |
|  | IGN<sup>32</sup> | ✓ | ✓ | 1.392 (0.020) | 1.094 (0.015) | 0.771 (0.009) | 0.762 (0.007) | 0.561 (0.019) | 0.786 (0.003) |
|  | SIGN<sup>6</sup> | ✓ | ✓ | **1.295 (0.010)** | **1.009 (0.015)** | **0.807 (0.004)** | **0.796 (0.003)** | **0.634 (0.01)** | **0.803 (0.002)** |
| Structure | SMINA<sup>33</sup> | ✓ | x | 2.078 (0.008) | 1.673 (0.011) | 0.609 (0.008) | 0.599 (0.007) | 0.342 (0.005) | 0.710 (0.003) |
|  | GNINA<sup>34</sup> | ✓ | x | 1.553 (0.017) | 1.232 (0.015) | 0.728 (0.009) | 0.737 (0.005) | 0.473 (0.005) | 0.771 (0.003) |
|  | dMaSIF<sup>35</sup> | ✓ | x | 1.528 (0.030) | 1.205 (0.037) | 0.720 (0.013) | 0.711 (0.015) | 0.507 (0.012) | 0.760 (0.007) |
|  | TankBind<sup>8</sup> | ✓ | x | 1.402 (0.021) | 1.085 (0.022) | 0.773 (0.008) | 0.764 (0.010) | 0.558 (0.022) | 0.788 (0.004) |
| Sequence | GraphDTA<sup>23</sup> | x | x | 1.568 (0.036) | 1.208 (0.028) | 0.698 (0.016) | 0.701 (0.012) | 0.475 (0.03) | 0.759 (0.005) |
|  | TransCPI<sup>3</sup> | x | x | 1.582 (0.034) | 1.243 (0.034) | 0.691 (0.022) | 0.674 (0.018) | 0.420 (0.031) | 0.741 (0.007) |
|  | MolTrans<sup>36</sup> | x | x | 1.616 (0.037) | 1.264 (0.022) | 0.675 (0.019) | 0.664 (0.019) | 0.443 (0.028) | 0.74 (0.008) |
|  | DrugBAN<sup>9</sup> | x | x | 1.436 (0.007) | 1.109 (0.005) | 0.762 (0.005) | 0.757 (0.004) | 0.540 (0.012) | 0.784 (0.003) |
|  | DGraphDTA<sup>4</sup> | x | x | 1.582 (0.034) | 1.243 (0.034) | 0.691 (0.022) | 0.674 (0.018) | 0.463 (0.028) | 0.747 (0.008) |
|  | WGNN-DTA<sup>17</sup> | x | x | 1.691 (0.020) | 1.332 (0.017) | 0.637 (0.014) | 0.614 (0.019) | 0.394 (0.025) | 0.721 (0.007) |
|  | STAMP-DPI<sup>18</sup> | x | x | 1.579 (0.026) | 1.197 (0.045) | 0.699 (0.020) | 0.680 (0.014) | 0.457 (0.025) | 0.747 (0.004) |
| PSICHIC |  | x | x | 1.336 (0.031) | 1.040 (0.029) | 0.792 (0.011) | 0.782 (0.013) | 0.594 (0.009) | 0.795 (0.006) |

|  | Methods | Hi-res<br>Structure | Co-crystal<br>Complex | PDBBind v2020 |  |  |  |  |  |
| --- | --- | --- | --- | --- | --- | --- | --- | --- | --- |
| | | | | RMSE ↓ | MAE ↓ | Pearson ↑ | Spearman ↑ | $r_m^2$ ↑ | CI ↑ |
| Complex | Pafnucy<sup>30</sup> | ✓ | ✓ | 1.435 (0.018) | 1.144 (0.018) | 0.635 (0.008) | 0.587 (0.008) | 0.348 (0.016) | 0.707 (0.004) |
|  | OnionNet<sup>31</sup> | ✓ | ✓ | 1.403 (0.012) | 1.103 (0.014) | 0.648 (0.007) | 0.602 (0.013) | 0.381 (0.011) | 0.717 (0.005) |
|  | IGN<sup>32</sup> | ✓ | ✓ | 1.404 (0.025) | 1.116 (0.030) | 0.662 (0.013) | 0.638 (0.021) | 0.385 (0.02) | 0.73 (0.009) |
|  | SIGN<sup>6</sup> | ✓ | ✓ | 1.373 (0.037) | 1.086 (0.030) | 0.685 (0.031) | 0.656 (0.044) | 0.398 (0.048) | 0.736 (0.02) |
| Structure | SMINA<sup>33</sup> | ✓ | x | 1.466 (0.008) | 1.161 (0.007) | 0.665 (0.005) | 0.663 (0.019) | 0.391 (0.031) | 0.74 (0.008) |
|  | GNINA<sup>34</sup> | ✓ | x | 1.740 (0.014) | 1.413 (0.015) | 0.495 (0.011) | 0.494 (0.011) | 0.209 (0.009) | 0.674 (0.004) |
|  | dMaSIF<sup>35</sup> | ✓ | x | 1.450 (0.032) | 1.136 (0.031) | 0.629 (0.018) | 0.588 (0.041) | 0.347 (0.029) | 0.71 (0.017) |
|  | TankBind<sup>8</sup> | ✓ | x | 1.345 (0.020) | 1.060 (0.021) | **0.718 (0.012)** | **0.689 (0.010)** | 0.404 (0.025) | 0.750 (0.006) |
| Sequence | GraphDTA<sup>23</sup> | x | x | 1.564 (0.063) | 1.223 (0.066) | 0.612 (0.016) | 0.570 (0.050) | 0.306 (0.039) | 0.703 (0.019) |
|  | TransCPI<sup>3</sup> | x | x | 1.493 (0.050) | 1.201 (0.037) | 0.604 (0.024) | 0.551 (0.029) | 0.255 (0.027) | 0.677 (0.011) |
|  | MolTrans<sup>36</sup> | x | x | 1.599 (0.060) | 1.271 (0.051) | 0.539 (0.057) | 0.474 (0.052) | 0.242 (0.045) | 0.666 (0.02) |
|  | DrugBAN<sup>9</sup> | x | x | 1.480 (0.046) | 1.159 (0.045) | 0.657 (0.018) | 0.612 (0.027) | 0.319 (0.021) | 0.72 (0.011) |
|  | DGraphDTA<sup>4</sup> | x | x | 1.493 (0.050) | 1.201 (0.037) | 0.604 (0.024) | 0.551 (0.029) | 0.312 (0.038) | 0.693 (0.011) |
|  | WGNN-DTA<sup>17</sup> | x | x | 1.501 (0.050) | 1.196 (0.055) | 0.605 (0.025) | 0.562 (0.028) | 0.311 (0.03) | 0.697 (0.01) |
|  | STAMP-DPI<sup>18</sup> | x | x | 1.503 (0.082) | 1.176 (0.067) | 0.653 (0.028) | 0.601 (0.027) | 0.327 (0.039) | 0.719 (0.011) |
| PSICHIC |  | x | x | **1.314 (0.049)** | **1.015 (0.031)** | 0.710 (0.027) | 0.686 (0.024) | **0.428 (0.047)** | **0.751 (0.009)** |

## References

1. Kitchen, D. B., Decornez, H., Furr, J. R. & Bajorath, J. Docking and scoring in virtual screening for drug discovery: methods and applications. Nat. Rev. Drug discovery 3, 935–949 (2004).

2. Hopkins, A. L. Predicting promiscuity. Nature 462, 167–168 (2009).

3. Chen, L. et al. Transformercpi: improving compound–protein interaction prediction by sequence-based deep learning with self-attention mechanism and label reversal experiments. Bioinformatics 36, 4406–4414 (2020).

4. Jiang, M. et al. Drug–target affinity prediction using graph neural network and contact maps. RSC Adv. 10, 20701–20712 (2020).

5. Bagherian, M. et al. Machine learning approaches and databases for prediction of drug–target interaction: a survey paper. Briefings Bioinforma. 22, 247–269 (2021).

6. Li, S. et al. Structure-aware interactive graph neural networks for the prediction of protein-ligand binding affinity. In Proceedings of the 27th ACM SIGKDD Conference on Knowledge Discovery & Data Mining, 975–985 (2021).

7. Dhakal, A., McKay, C., Tanner, J. J. & Cheng, J. Artificial intelligence in the prediction of protein–ligand interactions: recent advances and future directions. Briefings Bioinforma. 23, bbab476 (2022).

8. Lu, W. et al. Tankbind: Trigonometry-aware neural networks for drug-protein binding structure prediction. Adv. Neural Inf. Process. Syst. 35, 7236–7249 (2022).

9. Bai, P., Miljković, F., John, B. & Lu, H. Interpretable bilinear attention network with domain adaptation improves drug–target prediction. Nat. Mach. Intell. 1–11 (2023).

10. Ng, H. W. et al. Competitive molecular docking approach for predicting estrogen receptor subtype α agonists and antagonists. In BMC Bioinformatics, 1–15 (2014).

11. Rodríguez, D., Gao, Z.-G., Moss, S. M., Jacobson, K. A. & Carlsson, J. Molecular docking screening using agonist-bound gpcr structures: probing the a2a adenosine receptor. J. Chem. Inf. Model. 55, 550–563 (2015).

12. Kooistra, A. J., Leurs, R., De Esch, I. J. & de Graaf, C. Structure-based prediction of g-protein-coupled receptor ligand function: a β -adrenoceptor case study. J. Chem. Inf. Model. 55, 1045–1061 (2015).

13. Cai, T., Abbu, K. A., Liu, Y. & Xie, L. Deepreal: a deep learning powered multi-scale modeling framework for predicting out-of-distribution ligand-induced gpcr activity. Bioinformatics 38, 2561–2570 (2022).

14. Michel, M., Menéndez Hurtado, D. & Elofsson, A. Pconsc4: fast, accurate and hassle-free contact predictions. Bioinformatics 35, 2677–2679 (2019).

15. Rao, R., Meier, J., Sercu, T., Ovchinnikov, S. & Rives, A. Transformer protein language models are unsupervised structure learners. In International Conference on Learning Representations (2021).

16. Lin, Z. et al. Evolutionary-scale prediction of atomic-level protein structure with a language model. Science 379, 1123–1130 (2023).

17. Jiang, M. et al. Sequence-based drug-target affinity prediction using weighted graph neural networks. BMC Genomics 23 (2022).

18. Wang, P. et al. Structure-aware multimodal deep learning for drug–protein interaction prediction. J. Chem. Inf. Model. 62, 1308–1317 (2022).

19. Gainza, P. et al. Deciphering interaction fingerprints from protein molecular surfaces using geometric deep learning. Nat. Methods 17, 184–192 (2020).

20. Jumper, J. et al. Highly accurate protein structure prediction with alphafold. Nature 596, 583–589 (2021).

21. Wong, F. et al. Benchmarking alphafold-enabled molecular docking predictions for antibiotic discovery. Mol. Syst. Biol. 18, e11081 (2022).

22. He, X.-h. et al. Alphafold2 versus experimental structures: evaluation on g protein-coupled receptors. Acta Pharmacol. Sinica 44, 1–7 (2023).

23. Nguyen, T. et al. Graphdta: predicting drug–target binding affinity with graph neural networks. Bioinformatics 37, 1140–1147 (2021).

24. Corso, G., Stärk, H., Jing, B., Barzilay, R. & Jaakkola, T. S. Diffdock: Diffusion steps, twists, and turns for molecular docking. In The Eleventh International Conference on Learning Representations (2023).

25. Somnath, V. R., Bunne, C. & Krause, A. Multi-scale representation learning on proteins. Adv. Neural Inf. Process. Syst. 34, 25244–25255 (2021).

26. Corso, G., Cavalleri, L., Beaini, D., Liò, P. & Veličković, P. Principal neighbourhood aggregation for graph nets. Adv. Neural Inf. Process. Syst. 33, 13260–13271 (2020).

27. Liu, Z. et al. Pdb-wide collection of binding data: current status of the pdbbind database. Bioinformatics 31, 405–412 (2015).

28. Su, M. et al. Comparative assessment of scoring functions: the casf-2016 update. J. Chem. Inf. Model. 59, 895–913 (2018).

29. Stärk, H., Ganea, O., Pattanaik, L., Barzilay, R. & Jaakkola, T. Equibind: Geometric deep learning for drug binding structure prediction. In International Conference on Machine Learning, 20503–20521 (PMLR, 2022).

30. Stepniewska-Dziubinska, M. M., Zielenkiewicz, P. & Siedlecki, P. Development and evaluation of a deep learning model for protein–ligand binding affinity prediction. Bioinformatics 34, 3666–3674 (2018).

31. Zheng, L., Fan, J. & Mu, Y. Onionnet: a multiple-layer intermolecular-contact-based convolutional neural network for protein–ligand binding affinity prediction. ACS Omega 4, 15956–15965 (2019).

32. Jiang, D. et al. Interactiongraphnet: A novel and efficient deep graph representation learning framework for accurate protein–ligand interaction predictions. J. Medicinal Chem. 64, 18209–18232 (2021).

33. Koes, D. R., Baumgartner, M. P. & Camacho, C. J. Lessons learned in empirical scoring with smina from the csar 2011 benchmarking exercise. J. Chem. Inf. Model. 53, 1893–1904 (2013).

34. McNutt, A. T. et al. Gnina 1.0: molecular docking with deep learning. J. Cheminformatics 13, 1–20 (2021).

35. Sverrisson, F., Feydy, J., Correia, B. E. & Bronstein, M. M. Fast end-to-end learning on protein surfaces. In Proceedings of the IEEE/CVF Conference on Computer Vision and Pattern Recognition, 15272–15281 (2021).

36. Huang, K., Xiao, C., Glass, L. M. & Sun, J. Moltrans: molecular interaction transformer for drug–target interaction prediction. Bioinformatics 37, 830–836 (2021).

37. Zitnik, M., Sosicč, R., Maheshwari, S. & Leskovec, J. BioSNAP Datasets: Stanford biomedical network dataset collection (2018).

38. Liu, T., Lin, Y., Wen, X., Jorissen, R. N. & Gilson, M. K. Bindingdb: a web-accessible database of experimentally determined protein–ligand binding affinities. Nucleic Acids Res. 35, D198–D201 (2007).

39. Liu, H., Sun, J., Guan, J., Zheng, J. & Zhou, S. Improving compound–protein interaction prediction by building up highly credible negative samples. Bioinformatics 31, i221–i229 (2015).

40. Burley, S. K. et al. Rcsb protein data bank (rcsb. org): delivery of experimentally-determined pdb structures alongside one million computed structure models of proteins from artificial intelligence/machine learning. Nucleic Acids Res. 51, D488–D508 (2023).

41. Sun, J. et al. Excape-db: an integrated large scale dataset facilitating big data analysis in chemogenomics. J. Cheminformatics 9, 1–9 (2017).

42. Béquignon, O. J. et al. Papyrus: a large-scale curated dataset aimed at bioactivity predictions. J. Cheminformatics 15, 3 (2023).

43. Adasme, M. F. et al. Plip 2021: Expanding the scope of the protein–ligand interaction profiler to dna and rna. Nucleic Acids Res. 49, W530–W534 (2021).

44. Lin, H. et al. Discovery of potent and selective covalent protein arginine methyltransferase 5 (prmt5) inhibitors. ACS Medicinal Chem. Lett. 10, 1033–1038 (2019).

45. Rusere, L. N. et al. Hiv-1 protease inhibitors incorporating stereochemically defined p2 ligands to optimize hydrogen bonding in the substrate envelope. J. Medicinal Chem. 62, 8062–8079 (2019).

46. Yilmaz, N. K., Swanstrom, R. & Schiffer, C. A. Improving viral protease inhibitors to counter drug resistance. Trends Microbiol. 24, 547–557 (2016).

47. Draper-Joyce, C. J. et al. Structure of the adenosine-bound human adenosine a1 receptor–gi complex. Nature 558, 559–563 (2018).

48. Nguyen, A. T. et al. Extracellular loop 2 of the adenosine a1 receptor has a key role in orthosteric ligand affinity and agonist efficacy. Mol. Pharmacol. 90, 703–714 (2016).

49. Deng, Z., Chuaqui, C. & Singh, J. Structural interaction fingerprint (sift): a novel method for analyzing three-dimensional protein-ligand binding interactions. J. Medicinal Chem. 47, 337–344 (2004).

50. de Freitas, R. F. & Schapira, M. A systematic analysis of atomic protein–ligand interactions in the pdb. MedChemComm 8, 1970–1981 (2017).

51. Krivák, R. & Hoksza, D. P2rank: machine learning based tool for rapid and accurate prediction of ligand binding sites from protein structure. J. Cheminformatics 10, 1–12 (2018).

52. Bianchi, F. M., Grattarola, D. & Alippi, C. Spectral clustering with graph neural networks for graph pooling. In International Conference on Machine Learning, 874–883 (PMLR, 2020).

53. Rarey, M. & Dixon, J. S. Feature trees: a new molecular similarity measure based on tree matching. J. Comput. Mol. Des. 12, 471–490 (1998).

54. Jin, W., Yang, K., Barzilay, R. & Jaakkola, T. Learning multimodal graph-to-graph translation for molecule optimization. In International Conference on Learning Representations (2019).

55. Cai, T. et al. Graphnorm: A principled approach to accelerating graph neural network training. In International Conference on Machine Learning, 1204–1215 (PMLR, 2021).

56. Kingma, D. P. & Ba, J. Adam: A method for stochastic optimization. In International Conference on Learning Representations (2015).

57. Loshchilov, I. & Hutter, F. Decoupled weight decay regularization. In International Conference on Learning Representations (2019).

58. Steinegger, M. & Söding, J. Mmseqs2 enables sensitive protein sequence searching for the analysis of massive data sets. Nat. Biotechnol. 35, 1026–1028 (2017).

59. Khazanov, N. A. & Carlson, H. A. Exploring the composition of protein-ligand binding sites on a large scale. PLoS Comput. Biol. 9, e1003321 (2013).

60. Fukushima, K. Cognitron: A self-organizing multilayered neural network. Biol. Cybern. 20, 121–136 (1975).

61. Vaswani, A. et al. Attention is all you need. Adv. Neural Inf. Process. Syst. 30 (2017).

62. Baltos, J.-A. et al. Quantification of adenosine a1 receptor biased agonism: Implications for drug discovery. Biochem. pharmacology 99, 101–112 (2016).

63. Ismail-Fawaz, A. et al. An approach to multiple comparison benchmark evaluations that is stable under manipulation of the comparate set. arXiv preprint arXiv:2305.11921 (2023).

64. Clarivate. Cortellis drug discovery intelligence. https://www.cortellis.com/drugdiscovery/ (2023). Accessed: 02 02, 2023.

65. Bento, A. P. et al. An open source chemical structure curation pipeline using rdkit. J. Cheminformatics 12, 1–16 (2020).

66. Kumar, R. et al. Structure and energetics of ligand–fluorine interactions with galectin-3 backbone and side-chain amides: Insight into solvation effects and multipolar interactions. ChemMedChem 14, 1528–1536 (2019).

67. Kumar, R. et al. Substituted polyfluoroaryl interactions with an arginine side chain in galectin-3 are governed by steric-, desolvation and electronic conjugation effects. Org. & Biomol. Chem. 17, 1081–1089 (2019).

68. Rao, R. et al. Evaluating protein transfer learning with tape. Adv. Neural Inf. Process. Syst. 32 (2019).

69. Roy, K. et al. Some case studies on application of “rm2” metrics for judging quality of quantitative structure–activity relationship predictions: emphasis on scaling of response data. J. Comput. Chem. 34, 1071–1082 (2013).

