## Supplementary Information for "PSICHIC: physicochemical graph neural network for learning protein-ligand interaction fingerprints from sequence data"

#### Supplementary Methods

**1 PSICHIC Hyperparameters and Training Details 2**

**2 PDBBind Experimental Settings 2**

**3 PDB Resolution Sensitivity Analysis 5**

**4 Robustness Analysis via *In Silico* Structures 5**

**5 Benchmarking Performance of Sequence-based Methods 6**

**6 Protein-Ligand Interaction Response Dataset 8**

**7 Systematic Analysis of PSICHIC's Fingerprint Interpretability 8**

**8 Pharmacophore Analysis using PSICHIC's Fingerprints 8**

**9 PSICHIC for Virtual Screening Application 9**

**10 TankBind for Virtual Screening Application 10**

**11 PSICHIC Interpretability on $A_1$R-MIPS1 Interaction 10**

**12 PSICHIC Online Platform 11**

### 1 PSICHIC Hyperparameters and Training Details

We defined a hyperparameter search space as shown in Supplementary Table 1, and tuned PSICHIC on the PDBBind v2020 dataset. The best configuration was defined as the hyperparameter setting that achieved the lowest root-mean-square-error (RMSE) on the validation set. The same hyperparameters were used across all other experimental settings.

**Supplementary Table 1. PSICHIC Hyperparameter Search.** The hyperparameter options we searched through for PSICHIC on PDBBind v2020 validation set.

| PSICHIC Parameter | Search Space |
|---|---|
| Number of physicochemical convolutional layers | 2, **3**, 4 |
| Number of hidden dimensions, D | 50, 100, **200**, 400 |
| Number of Attention Heads | 1, **5**, 10 |
| Dropout Rate on Attention Logits | 0, **0.2**, 0.5 |
| Number of Protein Clustered Regions, K* | **[5, 10, 20]**, [10,15,20], [20, 20, 20] |
| Batch Size | 8, **16**, 32 |
| Non-linearities | **ReLU** |

* For 2 layers, we search [10,20] and [20,20]. For 4 layers, we search [5,10,15,20] and [20,20,20,20].

### 2 PDBBind Experimental Settings

The PDBBind database consists of experimentally derived protein-ligand complexes and their corresponding binding affinities. The availability of these complexes allowed us to comprehensively compare PSICHIC against state-of-the-art complex-based, structure-based, and sequence-based methods. In this database, the objective for computational methods is to predict the negative log-transformed binding affinity of protein-ligand pairs. For the scope of our paper, we specifically used two standard versions commonly cited in the literature: PDBBind v2016 and v2020. To compare PSICHIC, we executed 15 competing methods on the PDBBind v2016 and v2020 benchmarks five times and reported the average performances of these methods. We investigated a total of 4 complex-based approaches, 4 structure-based approaches, 3 two-dimensional sequence-based approaches, and 4 one-dimensional sequence-based approaches. Implementation details are provided below.

**Complex-Based Methods** rely on high-resolution, experimentally determined protein-ligand co-complex structures for making binding affinity predictions. These methods possess comprehensive knowledge of the protein-ligand complexes as model inputs. Each complex-based method employs a slightly different preprocessing pipeline, so when reproducing their performances on PDBBind v2016 and v2020, we followed the suggested pipeline from their original implementations.

- **Pafnucy** is a representative 3D Convolutional Neural Networks (CNN) method that models complex structures with a 3D grid. Pafnucy utilizes a 3D convolution to produce a feature map of the grid representation, treating the atoms of both proteins and ligands in the same manner. It requires high-resolution complex structures as the heavy atoms are discretized using a 3D grid with 1Å resolution. We preprocessed PDBBind datasets and reproduced results using the open-source code from https://gitlab.com/cheminfIBB/pafnucy.
- **OnionNet** creates two-dimensional interaction features by focusing on rotation-independent contacts between atom element pairs in complexes and utilizes to learn representations that can be used for making predictions. It requires high-resolution complex structures to obtain the dimensional interaction features. We preprocessed PDBBind datasets and reproduced results using the open-source code from https://github.com/zhenglz/onionnet.
- **IGN**, or Interaction Graph Net, models intra- and intermolecular forces by stacking two independent graph convolution modules were stacked to sequentially learn the intramolecular and intermolecular interactions. It, however, does not model the interdependent effects between intra- and intermolecular forces. To model intermolecular forces, IGN relies on co-complex structures to provide high-resolution atom-pair relations in the binding site. We preprocess PDBBind datasets and reproduce results using the open-source code from https://github.com/zjujdj/InteractionGraphNet/tree/master.
- **SIGN**, or Structure-aware Interactive Graph Network, composed of two primary elements: Polar-Inspired Graph Attention Layers (PGAL) and Pairwise Interactive Pooling (PiPool). PGAL models high-resolution spatial relations between the ligand atoms and protein pocket atoms, while PiPool learns to capture more distant, global interactions. We preprocessed PDBBind datasets and reproduced results using the open-source code from https://github.com/PaddlePaddle/PaddleHelix.

**Structure-Based Methods** employ high-resolution 3D protein structures and ligand SMILES strings as inputs for predicting binding affinity. Following the preprocessing pipeline from TankBind, we assumed that all structure-based methods have access to high-resolution experimentally-determined 3D protein structures, along with prior knowledge of protein chains located within 10 Å from the binding ligand:

- **TankBind** or Trigonometry-Aware Neural networK for BINDing structure prediction, incorporates a strong inductive bias through trigonometry constraints and focuses on all potential binding sites by dividing the protein into functional segments. The model uses innovative contrastive losses and local region negative sampling to optimize both binding interactions and affinity simultaneously. At the time of writing, TankBind is the current state-of-the-art for predicting protein-ligand binding affinity on PDBBind v2020, in terms of RMSE error. We preprocessed PDBBind datasets and reproduced results using the open-source code from `https://github.com/luwei0917/TankBind`.

- **dMaSIF** extends on MASIF to signficantly increase the computational efficiencies. Conceptually, MASIF proposes that the protein surface exhibits unique patterns of chemical and geometric characteristics that signify protein interactions with other biomolecules. By employing geodesic convolutions, it learns fingerprint vectors on protein surfaces. To benchmark its results on PDBBind datasets, we first concatenate the ligand and protein representations that are computed using three layers of PNA-GNN and dMASIF respectively. Then, we treat the concatenated representation as interaction fingerprint for a final feedforward network to predict the binding affinity values. We preprocessed PDBBind datasets and reproduced results using the open-source code from `https://github.com/FreyrS/dMaSIF`.

- **SMINA** enhances Autodock Vina by incorporating a novel scoring function and increased usability. The standard settings were employed, except for the alteration of –num_modes to 10. The search box was defined by utilizing the automatic box generation feature surrounding the receptor, along with the default 4Å buffer on all six sides. As the SMINA program does not provide a binding affinity value, we follow the original implementation of fitting a linear regression on the training data using AutoDock Vina features, namely Van Der Waals (vdw) forces, the charge-dependent desolvation term, the square of the number of internal torsions and the number of hydrophobic atoms. We preprocessed PDBBind datasets and reproduced results using the open-source code from `https://github.com/mwojcikowski/smina`.

- **QNINA** expands upon SMINA by incorporating a trained 3D CNN for evaluation purposes. With the exception of setting –num_modes to 10, default parameters were applied. The search box was established by employing the automatic box generation feature around the receptor, maintaining the standard 4Å buffer on each of the six sides. We preprocessed PDBBind datasets and reproduced results using the open-source code from `https://github.com/gnina/gnina`.

**2D Sequence-Based Methods** utilize ligand molecular graph and predicted protein contact map as inputs for predicting binding affinity. Consequently, these methods solely rely on the information derived from ligand SMILES and the complete protein sequence. Lacking prior knowledge of the binding pocket center, protein sequences extracted from the PDBBind datasets may extend up to 4,720 amino acids in length.

- **DGraphDTA** is a representative sequence-based method that leverages protein and ligand graphs in learning their respective representations using a graph neural network. It builds a ligand graph using molecular structure from RdKit, and a protein graph using a contact map predicted from Pconsc4. It concatenates both representations to obtain a fingerprint, implicitly modeling the interaction within the feature space. A feedforward neural network is then used to make accurate binding affinity predictions. We reproduced results by adapting the open-source code from `https://github.com/595693085/DGraphDTA`.

- **WGraphDTA** extends on DGraphDTA by obtaining a weighted contact map prediction using the ESM1 language model. Using ESM1 contact map prediction allows faster preprocessing of protein graphs as its contact map prediction is faster than Pcons4, and it can provide granular weighted edge representation between residues. To allow fair comparison between WGraphDTA and PSICHIC, we used ESM2 rather than ESM1 contact map prediction because it is more accurate. We also included embedding from the final layer of ESM2 as part of its residue node features. Hence, WGraphDTA may provide a direct comparison to PSICHIC, where WGraphDTA does not model protein-ligand interaction explicitly. We reproduced results by adapting the open-source code from `https://github.com/595693085/WGNN-DTA`.

- **STAMP-DPI** also leverages protein and ligand graphs in learning their respective representations using graph neural networks. It also explicitly models protein-ligand interaction using a Transformer model. Similar to WGraphDTA, we used ESM2 contact map prediction because it is more accurate and included embedding from the final layer of ESM2 as part of its residue node features. Hence, STAMP-DPI may provide a direct comparison to PSICHIC, where STAMP-DPI explicitly models protein-ligand interaction but without physicochemical constraints. We reproduced results by adapting the open-source code from `https://github.com/biomed-AI/STAMP-DPI`.

**1D Sequence-Based Methods** utilize ligand SMILES and protein sequences as inputs for predicting binding affinity. Similar to 2D sequence-based methods, these methods solely rely on the information derived from ligand SMILES and the complete protein sequence. Lacking prior knowledge of the binding pocket center, protein sequences extracted from the PDBBind datasets may extend up to 4,720 amino acids in length.

- **GraphDTA** is a representative sequence-based approach that utilizes a graph neural network to predict drug–target affinity. This method models ligands as graphs and utilizes graph neural networks to learn their representation. Concurrently, it employs a 1D CNN for modeling protein sequences and acquiring their representation. GraphDTA then concatenates both representations to obtain a fingerprint, implicitly modeling the interaction within the feature space. A feedforward neural network is then used to make accurate binding affinity predictions. We reproduced results using the open-source code from `https://github.com/thinng/GraphDTA`.

- **TransCPI** extends on GraphDTA by enriching the protein representation via Word2Vec and modeling the protein-ligand interaction using a Transformer architecture. It models the complete pairwise interaction between each atom-residue pair without any constraints to obtain a fingerprint for final interaction property prediction. As TransCPI is initially proposed for binary classification tasks only, we reproduced its results by swapping the cross entropy loss function into a mean-squared error loss using the open-source code from `https://github.com/lifanchen-simm/transformerCPI`.

- **MolTrans** is a protein-ligand interaction model that capitalizes on prior knowledge of ligand SMILES and protein amino acid sequence substructures. First, it employs the Frequent Consecutive Sub-sequence (FCS) algorithm to decompose ligand SMILES and protein sequence strings into their respective substructures. MolTrans then independently models ligand and protein representations on the substructures utilizing a dual Transformer architecture. Finally, MolTrans captures protein-ligand interactions by completely examining pairwise associations between all protein and ligand substructures to obtain a fingerprint for final interaction property prediction. We reproduced results using the open-source code from `https://github.com/kexinhuang12345/MolTrans`.

- **DrugBAN** also extends on GraphDTA by better modeling the protein-ligand interactions using bilinear attention networks. Firstly, it models the ligand molecular graph and protein sequence using separate graph neural networks and 1D-CNN blocks. Then, it employs a bilinear attention network to model the complete pairwise interaction between protein residue segments and ligand atoms to obtain a fingerprint for final interaction property prediction. As DrugBAN is initially proposed for binary classification tasks only, we reproduced its results by swapping the cross entropy loss function into a mean-squared error loss using the open-source code from `https://github.com/peizhenbai/DrugBAN`.

#### 3 PDB Resolution Sensitivity Analysis

We analyzed model sensitivity towards input quality of state-of-the-art sequence-based (PSICHIC), structure-based (TankBind), and complex-based (SIGN) methods using the PDBBind v2020 test set. We used Python package PyPDB to obtain the resolution quality of the complexes in the test set of PDBBind v2020. We failed to extract the resolution of one complex (PDB ID: 6V5L).

#### 4 Robustness Analysis via *In Silico* Structures

To compare the performance of TankBind and SIGN when only protein sequence data is available, we generated high-resolution 3D protein structures and co-complex structures using computational methods. TankBind requires 3D protein structures, while SIGN further necessitates 3D protein-ligand co-complex structures. To generate the 3D structures, we used AlphaFold and ESMFold. For co-complex structures, we further employed DiffDock to dock ligands to the 3D structures generated by AlphaFold and ESMFold. We selected DiffDock because it demonstrates state-of-the-art docking performance on the PDBBind v2020 dataset. However, given that our sequence input data could include up to 4,720 amino acid sequences, there is a possibility that AlphaFold, ESMFold, and/or DiffDock may fail to process this data. This limitation led us to conduct the comparison using a subset of the PDBBind v2020 test data, as detailed in Supplementary Table 2.

**Supplementary Table 2. Robustness Analysis via *In Silico* Structures.** Subset of PDBBind v2020 test set used to compare PSICHIC, TankBind and SIGN performances using sequence data only.

| | |
|---|---|
| **Comparison Subset** | 6N4E, 6JWA, 6GGB, 6PYB, 6I5P, 6HV2, 6OP9, 6D3Y, 6E3M, 6JIB, 6I64, 6JSN, 6PZ4, 6OY1, 6PY0, 6HOT, 6QLR, 6IC0, 6NRI, 6K3L, 6G2F, 6PGO, 6GZY, 6I8M, 6H7D, 6PKA, 6HZA, 6D5W, 6JAG, 5ZJY, 6I9A, 6E7M, 6MHB, 6JAQ, 6E3N, 6G2C, 6EFK, 6D3X, 6I77, 6JBE, 6G2E, 6N92, 6QMT, 6E3Z, 6E4V, 6GBW, 6I65, 6I78, 6I75, 6QLQ, 6HLE, 6O9B, 6G24, 6FFG, 6QGE, 6CYH, 6NPP, 6GGA, 6P8Z, 5ZK7, 6OIM, 6NRH, 5ZCU, 6OOZ, 6G2O, 6QLU, 6OY2, 6FFF, 6K1S, 6OI8, 6I67, 6D3Z, 6NP5, 6INZ, 6HHJ, 6OXU, 6AHS, 5ZK5, 6JSF, 6OIO, 6CJJ, 6OP0, 6HOQ, 6NV9, 6NP2, 6PNN, 5ZK9, 6A87, 6JSZ, 6MD6, 6JAP, 6N53, 6CKL, 6MHC, 6JON, 6MO9, 6OIQ, 6A6K, 6IBX, 6GGE, 6I63, 5ZLF, 6OXW, 6HZD, 6HLB, 6E4C, 6J9Y, 6N94, 6OXX, 6K2N, 6HBN, 6NW3, 6FTF, 6KJF, 6JAM, 6P8X, 6OIN, 6FFE, 6OVZ, 6A73, 6FCJ, 6IBY, 6JMF, 6OXR, 6PGP, 6J06, 6N4B, 6NP4, 6QMR, 6QLT, 6OXY, 6G27, 6HHH, 6JB0, 6I62, 6I61, 6H13, 6OOY, 6HHP, 6N55, 6CJP, 6J9W, 6N93, 6C85, 6OY0, 6MO8, 6CYG, 6E3O, 6H14, 6N97, 6N0M, 6NPM, 6M7H, 6MO0, 6QI7, 6G29, 6OM4, 6JAN, 6GDY, 6N8X, 6E6V, 6K04, 6I68, 6HLD, 6K05, 6OLX, 6I7A, 6OXZ, 6I76, 6HHI, 5ZE6, 6NXZ, 6HMT, 6IZQ, 6O9C, 6OXP, 6QLP, 6QLO, 6HHG, 6JAO, 6PNO, 6MO7, 6KQI, 6OXS, 6JAD, 6CJR, 6GGF, 6NPI, 6O3X, 6NY0, 6QLN, 6FE5, 6G2B, 6HHR, 6G9F, 6JSE, 6P8Y, 6GGD, 6H12, 6OIR, 6I41, 6NLJ, 6G3C, 5ZML, 5ZXK, 6I74, 6IBZ, 6N9L, 6KJD, 6QGF, 6HZB, 6ND3, 6JBB, 6O5U, 6G5U, 6OIE, 6D40, 6OXT, 6D07, 6NV7, 6O3Y, 6N96, 5ZR3, 6JSG, 6HOP, 6PNM, 6D08, 6HZC, 6CJS, 6NP3, 6JT3, 6HOU, 6I66, 6OIP, 6HMY, 6MHD, 6H9V, 6I8T, 6JB4, 6OD6, 6HOR, 6JID, 6QLS, 6OXV, 6OXQ |
| **PDB ID where AlphaFold, ESMFold and/or DiffDock fails to generate complexes** | 6QUW, 6QUV, 6GJ5, 6UHU, 6G25, 6V5L, 6QTQ, 6RNU, 6UVP, 6QQT, 6QWI, 6QTX, 6UFO, 6QRF, 6QTO, 6QTR, 6GJ6, 6UEG, 6QQW, 6SEO, 6QTS, 6QRE, 6QRC, 6QQV, 6QRG, 6QR1, 6R4K, 6S56, 6S57, 6QQQ, 6UVV, 6QTM, 6SFC, 6EEB, 6QQU, 6OS5, 6OTT, 6QR2, 6QTW, 6SEN, 6UFN, 6O5G, 6QXA, 6GJ8, 6OS6, 6UVY, 6QXD, 6GJ7, 6QRA, 6S55, 6QR4, 6ROT, 6Q38, 6BQD, 6QW8, 6S9X, 6UII, 6E5S, 6QR3, 6TE6, 6RTN, 6MO2, 6E13, 6T6A, 6QR9, 6QQZ, 6UIM, 6PYA, 6Q4Q, 6QR0, 6UHV, 6QRD, 6ST3, 6MOA, 6QFE, 6UIL, 6QR7, 6QSZ, 6RR0, 6S9W, 6NRG, 6MJI, 5ZJZ, 6GWE, 6S07, 6NRJ, 6UWP, 6Q36, 6MJ4, 6MIY, 6NRF, 6NSV, 6MJJ, 6MJA, 6MJQ, 6DQL, 6CF7, 6UWV, 6E6J, 6A1C, 6MIV, 6O0H, 6AGT, 6QZH, 6RPG, 6RZ6, 6R7D, 6JUT, 6E3P, 6NT2, 6DZ2, 6DZ3, 6DZ0, 6DYZ, 6IQL, 6TEN, 6E6W, 6TEL |

#### 5 Benchmarking Performance of Sequence-based Methods

To evaluate the performance of PSICHIC and leading sequence-based methods across a diverse range of drug discovery settings, we partitioned the datasets (Human, BioSNAP, and BindingDB) used in DrugBAN into training, validation, and test sets using a 70/10/20 split, with 70% of the data allocated for training, 10% for validation, and 20% for testing. We then used three different strategies for this partitioning: random split, unseen ligand scaffold split, and unseen protein target split. In the random split strategy, protein-ligand interaction data were randomly assigned to the training, validation, and test sets. For the unseen ligand scaffold split, unique ligand scaffolds were allocated to the training, validation, and test sets in such a way that there was no overlap between the scaffolds. Similarly, in the unseen protein target split setting, unique protein targets were assigned to the training, validation, and test sets without any overlap. Under these latter two conditions, models could not rely on known ligand or protein features present in the training data to make predictions.

We reproduced the results of all sequence-based methods in a manner similar to the original implementations of these methods on the PDBBind datasets. The key difference was that the final layer was a classification head, and the methods were optimized using binary cross-entropy loss. For each setting, we ran all methods five times and reported their average Area Under the Receiver Operating Characteristics Curve (AUC-ROC) and Area Under the Precision-Recall Curve (AUC-PRC) performance in Supplementary Table 3 for Human, BioSNAP, and BindingDB datasets, respectively. We performed all pairwise comparisons across the sequence-based methods in Supplementary Table 4, with the top three methods being PSICHIC, DrugBAN, and STAMP-DPI, in that order. These three methods were subsequently compared in the main text sections.

**Supplementary Table 3. Benchmarking Binary Classification Across Nine Different Drug Discovery Scenarios**. The drug discovery scenarios were generated using three different data-splitting strategies (random split, unseen ligand split, and unseen protein split) applied to three datasets: Human, BioSNAP, and BindingDB.

| | **Methods** | **Random Split** | | **Unseen Ligand Scaffold** | | **Unseen Protein Target** | |
|---|---|---|---|---|---|---|---|
| | | **AUC-ROC** | **AUC-PRC** | **AUC-ROC** | **AUC-PRC** | **AUC-ROC** | **AUC-PRC** |
| **Human** | GraphDTA | 0.969 (0.004) | 0.960 (0.007) | 0.940 (0.005) | 0.926 (0.007) | 0.901 (0.012) | 0.907 (0.015) |
| | TransCPI | 0.951 (0.008) | 0.944 (0.01) | 0.937 (0.004) | 0.926 (0.008) | 0.866 (0.032) | 0.912 (0.008) |
| | MolTrans | 0.979 (0.002) | 0.974 (0.004) | 0.912 (0.004) | 0.903 (0.009) | 0.917 (0.015) | 0.925 (0.013) |
| | DrugBAN | 0.981 (0.002) | 0.971 (0.009) | 0.944 (0.004) | **0.937 (0.005)** | 0.923 (0.007) | 0.93 (0.009) |
| | DGraphDTA | 0.967 (0.003) | 0.963 (0.005) | 0.877 (0.005) | 0.857 (0.018) | **0.953 (0.003)** | **0.959 (0.002)** |
| | WGNN-DTA | 0.970 (0.004) | 0.967 (0.005) | 0.881 (0.009) | 0.855 (0.027) | 0.951 (0.011) | 0.958 (0.009) |
| | STAMP-DPI | 0.962 (0.003) | 0.954 (0.003) | 0.907 (0.008) | 0.893 (0.009) | 0.854 (0.002) | 0.881 (0.009) |
| | **PSICHIC** | **0.983 (0.002)** | **0.979 (0.004)** | **0.945 (0.004)** | 0.930 (0.008) | 0.943 (0.007) | 0.950 (0.009) |
| **BioSNAP** | GraphDTA | 0.887 (0.003) | 0.878 (0.004) | 0.878 (0.002) | 0.876 (0.006) | 0.741 (0.008) | 0.795 (0.007) |
| | TransCPI | 0.892 (0.002) | 0.894 (0.004) | 0.876 (0.005) | 0.873 (0.006) | 0.754 (0.016) | 0.809 (0.010) |
| | MolTrans | 0.895 (0.005) | 0.888 (0.017) | 0.861 (0.004) | 0.860 (0.006) | 0.693 (0.010) | 0.745 (0.007) |
| | DrugBAN | 0.906 (0.002) | 0.912 (0.003) | 0.888 (0.006) | 0.892 (0.005) | 0.712 (0.015) | 0.772 (0.015) |
| | DGraphDTA | 0.869 (0.008) | 0.886 (0.008) | 0.860 (0.004) | 0.877 (0.003) | 0.771 (0.016) | 0.830 (0.013) |
| | WGNN-DTA | 0.855 (0.035) | 0.852 (0.054) | 0.849 (0.016) | 0.867 (0.013) | 0.786 (0.012) | 0.836 (0.010) |
| | STAMP-DPI | 0.895 (0.003) | 0.900 (0.005) | 0.886 (0.004) | 0.882 (0.008) | 0.821 (0.009) | 0.832 (0.065) |
| | **PSICHIC** | **0.920 (0.004)** | **0.918 (0.003)** | **0.895 (0.005)** | **0.896 (0.006)** | **0.876 (0.008)** | **0.904 (0.006)** |
| **BindingDB** | GraphDTA | 0.936 (0.002) | 0.912 (0.002) | 0.917 (0.003) | 0.884 (0.005) | 0.629 (0.014) | 0.501 (0.027) |
| | TransCPI | 0.951 (0.002) | 0.937 (0.002) | 0.923 (0.003) | 0.897 (0.005) | 0.682 (0.035) | 0.589 (0.043) |
| | MolTrans | 0.953 (0.002) | 0.936 (0.003) | 0.923 (0.003) | 0.891 (0.011) | 0.573 (0.048) | 0.483 (0.052) |
| | DrugBAN | **0.961 (0.002)** | **0.949 (0.005)** | **0.939 (0.001)** | **0.914 (0.003)** | 0.635 (0.010) | 0.523 (0.005) |
| | DGraphDTA | 0.896 (0.011) | 0.863 (0.019) | 0.884 (0.006) | 0.842 (0.011) | 0.623 (0.024) | 0.516 (0.045) |
| | WGNN-DTA | 0.896 (0.008) | 0.861 (0.012) | 0.898 (0.014) | 0.862 (0.022) | 0.639 (0.067) | 0.521 (0.075) |
| | STAMP-DPI | 0.927 (0.003) | 0.905 (0.003) | 0.906 (0.004) | 0.870 (0.004) | 0.675 (0.005) | 0.552 (0.017) |
| | **PSICHIC** | 0.948 (0.002) | 0.925 (0.007) | 0.923 (0.002) | 0.887 (0.008) | **0.728 (0.019)** | **0.603 (0.021)** |

* The **best** and second-best performances are marked in bold and underlined, respectively.

| Mean-ROC-AUC | PSICHIC<br>0.9068 | DrugBAN<br>0.8766 | STAMP-DPI<br>0.8703 | TransCPI<br>0.8702 | GraphDTA<br>0.8692 | WGNN-DTA<br>0.8583 | MolTrans<br>0.8563 | DGraphDTA<br>0.8556 |
|---|---|---|---|---|---|---|---|---|
| PSICHIC<br>0.9068 | Mean-Difference<br>r>c / r=c / r<c<br>Wilcoxon p-value | 0.0302<br>7 / 0 / 2<br>0.1641 | **0.0364**<br>**9 / 0 / 0**<br>**0.0039** | **0.0366**<br>**7 / 1 / 1**<br>**0.0176** | **0.0376**<br>**9 / 0 / 0**<br>**0.0039** | **0.0484**<br>**8 / 0 / 1**<br>**0.0078** | **0.0504**<br>**7 / 1 / 1**<br>**0.0241** | **0.0512**<br>**8 / 0 / 1**<br>**0.0078** |
| DrugBAN<br>0.8766 | -0.0302<br>2 / 0 / 7<br>0.1641 | - | 0.0062<br>7 / 0 / 2<br>0.4961 | 0.0063<br>7 / 0 / 2<br>0.4258 | 0.0074<br>7 / 0 / 2<br>0.1641 | 0.0182<br>6 / 0 / 3<br>0.3008 | **0.0202**<br>**9 / 0 / 0**<br>**0.0039** | 0.0210<br>7 / 0 / 2<br>0.2031 |
| STAMP-DPI<br>0.8703 | **-0.0364**<br>**0 / 0 / 9**<br>**0.0039** | -0.0062<br>2 / 0 / 7<br>0.4961 | - | 0.0001<br>4 / 0 / 5<br>0.5703 | 0.0011<br>4 / 0 / 5<br>0.9102 | 0.0120<br>7 / 0 / 2<br>0.2031 | 0.0140<br>3 / 1 / 5<br>1.0000 | 0.0148<br>7 / 0 / 2<br>0.1641 |
| TransCPI<br>0.8702 | **-0.0366**<br>**1 / 1 / 7**<br>**0.0176** | -0.0063<br>2 / 0 / 7<br>0.4258 | -0.0001<br>5 / 0 / 4<br>0.5703 | - | 0.0010<br>5 / 0 / 4<br>0.7344 | 0.0119<br>6 / 0 / 3<br>0.3594 | 0.0139<br>4 / 1 / 4<br>0.6350 | 0.0147<br>6 / 0 / 3<br>0.3594 |
| GraphDTA<br>0.8692 | **-0.0376**<br>**0 / 0 / 9**<br>**0.0039** | -0.0074<br>2 / 0 / 7<br>0.1641 | -0.0011<br>5 / 0 / 4<br>0.9102 | -0.0010<br>4 / 0 / 5<br>0.7344 | - | 0.0109<br>5 / 0 / 4<br>0.4258 | 0.0129<br>5 / 0 / 4<br>0.3008 | 0.0136<br>7 / 0 / 2<br>0.2031 |
| WGNN-DTA<br>0.8583 | **-0.0484**<br>**1 / 0 / 8**<br>**0.0078** | -0.0182<br>3 / 0 / 6<br>0.3008 | -0.0120<br>2 / 0 / 7<br>0.1641 | -0.0119<br>3 / 0 / 6<br>0.3594 | -0.0109<br>4 / 0 / 5<br>0.4258 | - | 0.0020<br>3 / 0 / 6<br>1.0000 | 0.0028<br>5 / 1 / 3<br>0.3127 |
| MolTrans<br>0.8563 | **-0.0504**<br>**1 / 1 / 7**<br>**0.0241** | **-0.0202**<br>**0 / 0 / 9**<br>**0.0039** | -0.0140<br>5 / 1 / 3<br>1.0000 | -0.0139<br>4 / 1 / 4<br>0.6350 | -0.0129<br>4 / 0 / 5<br>0.3008 | -0.0020<br>6 / 0 / 3<br>1.0000 | - | 0.0008<br>6 / 0 / 3<br>0.9102 |
| DGraphDTA<br>0.8556 | **-0.0512**<br>**1 / 0 / 8**<br>**0.0078** | -0.0210<br>2 / 0 / 7<br>0.2031 | -0.0148<br>2 / 0 / 7<br>0.1641 | -0.0147<br>3 / 0 / 6<br>0.3008 | -0.0136<br>2 / 0 / 7<br>0.2031 | -0.0028<br>3 / 1 / 5<br>0.3127 | -0.0008<br>3 / 0 / 6<br>0.9102 | **If in bold, then**<br>**p-value < 0.05** |

Mean-Difference

0.05 0 −0.05

**Supplementary Table 4. Multi-Comparison Matrix for Benchmarking Sequence-based Methods**. The matrix displays pairwise comparisons among PSICHIC, DrugBAN, STAMP-DPI, TransCPI, GraphDTA, WGNN-DTA, MolTrans, and DGraphDTA across 9 drug discovery settings from Human, BioSNAP, and BindingDB benchmark datasets. The evaluations are made using random split, unseen ligand scaffold split, and unseen protein target split. Performance is assessed using the Area Under the Receiver Operating Characteristics Curve (ROC-AUC). The colors on the Heat Map signify the mean differences in ROC-AUC performance. A positive difference, indicated in red, signifies that the method in the row outperforms the method in the column on average. Each cell contains three lines detailing performance metrics: the first line shows the average difference in ROC-AUC between the method in the row and the method in the column; the second line shows the number of wins/ties/losses across various drug discovery settings; and the third line indicates the p-value, determined through the Wilcoxon Signed Rank Test. Text within each cell is presented in **BOLD** if the p-value is lower than 0.05.

#### 6 Protein-Ligand Interaction Response Dataset

As a data-driven framework, PSICHIC is capable of learning to predict any interaction properties from a labeled sequence dataset. To equip PSICHIC with the ability to forecast the type of response a protein will elicit upon binding to a ligand, we manually collected and curated response-type data from high-quality databases. This curation process is illustrated in Extended Data Fig. 3.

We initially collected high-quality functional effect data from the Cortellis Drug Discovery database, setting the filter to "Receptor" and including only proteins with more than 20 data samples. This data was downloaded on February 2, 2023. Following this manual curation process, we obtained 22,085 protein-ligand pairs labeled as agonists and 17,211 labeled as antagonists. Each sample in the dataset represents a protein-ligand sequence pair and specifies whether the ligand is an agonist or an antagonist.

However, the Cortellis dataset lacked negative data or decoys, which are essential for model training. Instead of heuristically generating decoys, we chose a more rigorous approach. We utilized the two largest publicly available datasets, ExCAPE-ML (with potency values denoted as pXC50) and Papyrus (with binding affinity values noted as pKi/pKd), to extract all protein-ligand pairs with values below 5, labeling them as non-binders. For the ExCAPE-ML dataset, we exclusively selected protein-ligand pairs with available pXC50 values. Similarly, in the Papyrus dataset, we strictly focused on protein-ligand pairs designated as high-quality.

Finally, as all three databases (Cortellis Drug Discovery, ExCAPE-ML, and Papyrus) originate from different sources, we needed to standardize them. To do this, we employed the ChEMBL pipeline for data standardization, normalization, and amalgamation. In this way, we obtained a comprehensively curated dataset comprising 160,910 unique protein-ligand pairs: 22,085 agonists, 17,211 antagonists, and 121,614 non-binders. Our dataset includes 131 unique protein receptors and 128,122 unique ligands. Alongside this study, we are also making this highly curated dataset publicly available.

The curated functional effect dataset also includes potency values for agonists (EC50) and antagonists (IC50). However, because these potency values depend on the specific assays used, we chose to omit them from the training process. Instead, we utilized these values for the UMAP plot, as shown in Fig. 2 of the main results section.

#### 7 Systematic Analysis of PSICHIC's Fingerprint Interpretability

This section outlines the experimental details used to analyze the interpretability of PSICHIC, as discussed in the main section of the paper.

To investigate the interpretability of PSICHIC, we extracted all complexes from the PDBBind v2020 test set. These complexes consist of protein-ligand pairs that were not encountered during training and feature co-crystallized structures for both visualization and systematic evaluation. This approach enables us to assess how well PSICHIC performs on data it has not previously seen. Our analysis of PSICHIC's interpretability with respect to proteins and ligands is conducted as follows:

- **Protein:** Proximity of Residue to Binding Site *versus* PSICHIC Residue Importance Score.
- **Ligand:** Presence of Atom Forming Interactions with Binding Residues *versus* PSICHIC Atom Importance Score.

To assess the proximity of a residue to the binding site, we calculated this value as 1 minus normalized distance between the residue and the binding site. This way, the proximity of a residue falls within the range of 0 to 1. The distance was defined as the minimum Euclidean distance between any atom of the residue and any atom within the entire ligand. We then compared this proximity score against the median value of PSICHIC Residue Importance Scores, obtained from five runs on PDBBind v2020, using Pearson Correlation for the comparison.

To assess whether an atom in a ligand interacts with any binding residues, we employed the Protein-Ligand Interaction Profiler (PLIP). PLIP identifies various types of interactions, including hydrophobic interactions, hydrogen bonds, water bridges, salt bridges, metal complexes, pi-stacking, pi-cation, and halogen bonds. If an atom engages in multiple types of interactions, all applicable labels were included. Finally, we compared the average importance scores assigned to these labeled ligand atoms, categorizing them based on their type of interaction with binding residues. These results are depicted in Fig. 3 of the main text. We excluded halogen bond interactions from the analysis, as the dataset contains only 20 such interactions. However, PSICHIC, on average, assigned the highest importance scores (0.765) to atoms involved in halogen bonds with binding residues.

#### 8 Pharmacophore Analysis using PSICHIC's Fingerprints

To demonstrate the applicability of PSICHIC in pharmacophore analysis, we utilized 14 complex structures of Galectin-3 co-bound with different ligands from the PDBBind v2020 test set, as provided in two separate studies.

Before performing pharmacophore analysis on the ligands, we examined the Pearson correlation of PSICHIC residue importance scores in relation to the proximity of each residue to its binding site, as well as its confidence score. The latter is computed using the Gini coefficient (Extended Data Fig. 4a). Consistent with our analysis in the main text, PSICHIC assigned higher confidence scores (i.e., assigned higher importance scores to fewer residues) when it predicted the binding affinity to be higher. Moreover, the scores showed better correlation with the proximity of residues to the binding site. As an illustration, the importance scores for residues in 6QLS are dispersed around the protein (as seen in the upper left of Extended Data Fig. 4b), even extending to the opposite side of the protein or 180 degrees away from the binding site region (as seen in the lower left of Extended Data Fig. 4b). This occurred despite the highest importance being attributed to the binding residues. In contrast, 6I76, whose ligand was predicted by PSICHIC to have the highest affinity, was assigned importance scores that are predominantly focused on the binding sites (as seen in the upper right of Extended Data Fig. 4b). This observation affirmed that PSICHIC adheres to physicochemical principles that dictate protein-ligand interactions, and determines the protein binding site based on the ligand-induced interactions rather than merely memorizing the protein features.

In our pharmacophore analysis of binding ligands, PSICHIC displayed a unique capability of identifying various functional groups crucial for forming the binding interaction with Galectin-3C (Extended Data Fig. 4c). Across all cases, PSICHIC attributed significant importance to the fluorine atoms in the ligands. This finding aligned closely with original studies suggesting that due to the presence of several suitably oriented main-chain and side-chain amide groups, the pocket in Galectin-3C served as an exemplary model for investigating fluorine–amide interactions.

For comparative purposes, PSICHIC did not universally assign the highest importance to fluorine atoms (Extended Data Fig. 4d). The first three small-molecule ligands (PDB ID: 6OIN, 6OIQ, and 6OIR) shows three inhibitors of histone acetyltransferase, KAT6A. In these examples, PSICHIC did not highlight fluorine atoms as the most significant. Instead, PSICHIC identified the sulfonyl group, which forms hydrogen bonds with the binding residues of the Histone Acetyltransferase, as having the highest importance. In another example with the small-molecule ligand, 6TEN, PSICHIC assigned a higher importance score to the chlorine atom of chloranylpyridin due to its pi-stacking interaction with the binding residues. The fluorine atoms in this instance do not form any specific interactions with the binding residues, thus further illustrating that PSICHIC did not prioritize fluorine when the fluorine atom does not form important interaction with the target protein.

#### 9 PSICHIC for Virtual Screening Application

As a data-driven framework, PSICHIC learns to predict interaction properties from labeled sequence datasets. To enhance PSICHIC's capacity for predicting both the binding affinity of a protein-ligand pair and the functional effect a protein will have upon binding to a ligand, we manually collected and curated functional effect type data from large-scale databases. The scale of this database ensured that PSICHIC could capitalize on large-scale sequence-only databases to navigate the vast protein-ligand interaction landscape and effectively screen the library, rather than relying on strenuous extrapolation by PSICHIC itself. The process is illustrated in Extended Data Fig. 6.

From the Protein-Ligand Functional Effect dataset, we extracted all samples, each consisting of a protein, a ligand, and a label indicating whether the ligand is an agonist, antagonist or non-binder. From ExCAPE-ML, we extracted all protein-ligand pairs with a potency lower than 5 in terms of pXC50 and labeled them as non-binders, and those greater than 7 in terms of pXC50 were labeled as binders. From Papyrus, we extracted all protein-ligand pairs with high-quality binding affinity labels. Similarly, pairs with binding affinity lower than 5 in terms of pKi/Kd were labeled as non-binders, and those greater than 7 in terms of pKi/Kd were labeled as binders. We then obtained a Large-Scale Dataset by standardizing, normalizing, and combining all three databases, resulting in a dataset with approximately 3 million unique protein-ligand pairs labeled with binding affinity and/or response type labels. As all three databases (Protein-Ligand Functional Effect, ExCAPE-ML, and Papyrus) originated from different sources and required standardization, we used ChEMBL pipeline to standardize and normalize the molecular ligand data. After standardization, we obtained 618,247 completely labeled protein-ligand pairs with binding affinity and response type labels, as well as 2,341,057 partially labeled pairs. The large-scale dataset comprehensively includes a total of 5,107 unique proteins and 1,084,834 unique ligands.

While the large-scale interaction dataset is diverse, the number of samples per protein, given the interaction types, is extremely imbalanced. Specifically, the top 50 proteins (or 1%) out of 5,107 account for more than half of the dataset (see Extended Data Fig. 8a). Furthermore, most protein-ligand interactions are characterized by non-binders. If PSICHIC were to be trained on this large-scale dataset using random batch sampling, it would likely overfit to a handful of proteins or response types, undermining the purpose of training on a large-scale dataset. This issue is not resolved, and may even be exacerbated, if we group each protein and ligand scaffold together and sample each group equally (see Extended Data Fig. 8b). Therefore, this situation necessitated the development of a better training strategy that effectively balances the uniqueness of protein-ligand interaction samples, capturing the diversity of proteins, ligands, and interaction types. To ensure that PSICHIC could effectively train and learn from the large-scale dataset, the optimization process involved addressing two issues: (i) designing a loss function that can handle samples with partial labels, and (ii) developing a batch sampling strategy that ensures PSICHIC can learn from a diverse range of protein-ligand interactions and does not overfit to certain types of biases (i.e., imbalance in number of samples for each protein).

To address the partial label issues in the large-scale dataset, we designed a multi-task loss function capable of handling these training pairs. Partial annotation occurs in two instances: (i) when we possess the response class type (i.e., agonist, antagonist, or non-binder) but lack the binding affinity label, and (ii) when we have the binding affinity labels indicating strong binding with the protein (i.e., the ligand is a binder), but lack knowledge about the response type. In the first case (i), we bypass the computation of loss and calculate only the cross-entropy loss for the prediction of the response type. In the second case (ii), we aim for the model to predict the probability of the response type being a non-binder as low as possible. Therefore, we minimize the loss function as $-log(p \neq \text{non-binder})$ or $-(log(p_{agonist}) + log(p_{antagonist}))$. In this manner, we compute the cross-entropy loss normally when we know the exact label. However, when we do not know whether it is agonist or antagonist, we maximize the probability that it is *not* a non-binder. During batch training of PSICHIC, this did not cause instability as we took the average loss across a batch size, $B$ of 16, and each batch likely contained at least one label for either objective. Collectively, the supervision loss, $\mathbf{O}_{\text{APPLICATION(S)}}$ for each batch is taken as the sum over the batch average of $\mathbf{O}_{\text{MSE}}$ and $\mathbf{O}_{\text{CE}}$: $\frac{1}{|\text{S}_{\text{aff}}|}\sum_{i=1}^{B}\mathbf{O}_{\text{MSE}_i} + \frac{1}{|\text{S}_{\text{res}}|}\sum_{i=1}^{B}\mathbf{O}_{\text{CE}_i}$ where $|\text{S}_{\text{aff}}|$ and $|\text{S}_{\text{res}}|$ are the number of samples for affinity values and response class labels respectively. The multi-objective training process is depicted in Extended Data Fig. 7.

To address the imbalanced nature of the large-scale dataset, we adopted a pretrain-then-finetune strategy for training PSICHIC. In both the pretraining and finetuning steps, we carefully sampled the protein-ligand pairs to balance the uniqueness of protein-ligand interaction samples, capturing the diversity of proteins, ligands, and interaction types (see Extended Data Fig. 8). This will be achieved by assigning a probability weight to each protein-ligand pair, indicating how likely the pair will be sampled during training. The sampling weights in both stages were assigned as follows:

- **Pre-training**: We categorized each protein-ligand interaction sample based on its protein and whether it is a binder or a non-binder. For example, we created groups like ProteinA_Binder, ProteinA_Non-Binder, ..., ProteinZ_Binder, ProteinZ_Non-Binder. We then assigned a sampling weight to each group proportional to the square root of the group's size, capping this at the 90th percentile to prevent overrepresentation. Within each group, we ensured that each ligand scaffold is sampled equally. The resulting sampling distribution by protein and response type is illustrated in Extended Data Fig. 8. However, as illustrated in the right pie chart of Extended Data Fig. 8, even after adjusting for the sampling weights, the agonist and antagonist class types remain sparsely sampled. An alternative approach would be to increase the sampling of agonist and antagonist classes. However, this could lead to their over-representation within PSICHIC's pretraining process. This, in turn, could significantly reduce the diversity of the training data samples. Such an outcome would be in direct conflict with the intention of training on a large-scale dataset, which aimed to explore a comprehensive interaction space. As a resolution, we relied on fine-tuning to address this issue.

- **Fine-tuning**: The second stage, which involved fine-tuning PSICHIC for in silico screening of the adenosine A1 receptor protein, commenced by training PSICHIC for only 1,000 iterations. During this fine-tuning process, we *only included agonist and antagonist interactions* for $A_1R$ and any $A_1R$-related proteins. $A_1R$-related protein is defined as any protein with a sequence overlap greater than 50%, as defined by MMSeq. This approach ensured that PSICHIC was better adapted for the virtual screening of specific protein targets.

#### 10 TankBind for Virtual Screening Application

To screen our in-house MIPS library using TankBind, the leading structure-based method, we needed a 3D protein structure. For this purpose, we selected the experimentally-determined structure of the human adenosine $A_1$ receptor–Gi complex in its functionally active state. This structure is available in the Protein Data Bank under the PDB ID 6D9H. Following the TankBind open-source workflow, we first isolated only the Adenosine $A_1$ receptor ($A_1R$) from this complex structure. We then employed the publicly available TankBind model and its weights to screen the MIPS library. The binding affinity values predicted by TankBind served as the ranking measure for potential $A_1R$ agonists.

#### 11 PSICHIC Interpretability on $A_1R$-MIPS1 Interaction

To visualize PSICHIC's interpretation of the protein residues interacting with the sole agonist compound (MIPS1), we used the active $A_1R$ structure from PDB ID: 6D9H. However, the sequence information utilized by PSICHIC came from the UniProt database for the Human Adenosine A1 Receptor, which did not perfectly align with the residues in the PDB $A_1R$ structure. To reconcile this, we first aligned the two sequences and mapped the corresponding residue scores from the UniProt sequence to the residues in the PDB $A_1R$ structure. Due to some residue scores being omitted because of the imperfect alignment, we visualized the interpretability of protein-residue interactions in Fig. 4c of main text by coloring the residues according to their ranking percentile. A higher (or lower) ranking was represented by a darker (or lighter) red hue.

#### 12 PSICHIC Online Platform

The PSICHIC Virtual Screening Platform, operating on Google Colaboratory, offers a specialized approach to predicting protein-ligand interactions. Utilizing only sequence data, PSICHIC achieves high accuracy in binding affinity predictions and can effectively predict the functional effects of a ligand on its target protein. The platform also generates interpretable fingerprints, identifying key residues in the protein and atoms in the ligand involved in the interaction. Notably, PSICHIC can perform these analyses while screening up to 100,000 compounds per hour, facilitating pharmacophore and targeted mutagenesis studies. With its requirement for only a CSV file containing Protein Sequence and Ligand SMILES pairs, PSICHIC is both accessible and efficient (see Supplementary Fig. 1). Its user-friendly, interactive platform democratizes access to this specialized method, aiming to advance the field of virtual screening and deepen our understanding of protein-ligand interactions (see Supplementary Fig. 2).

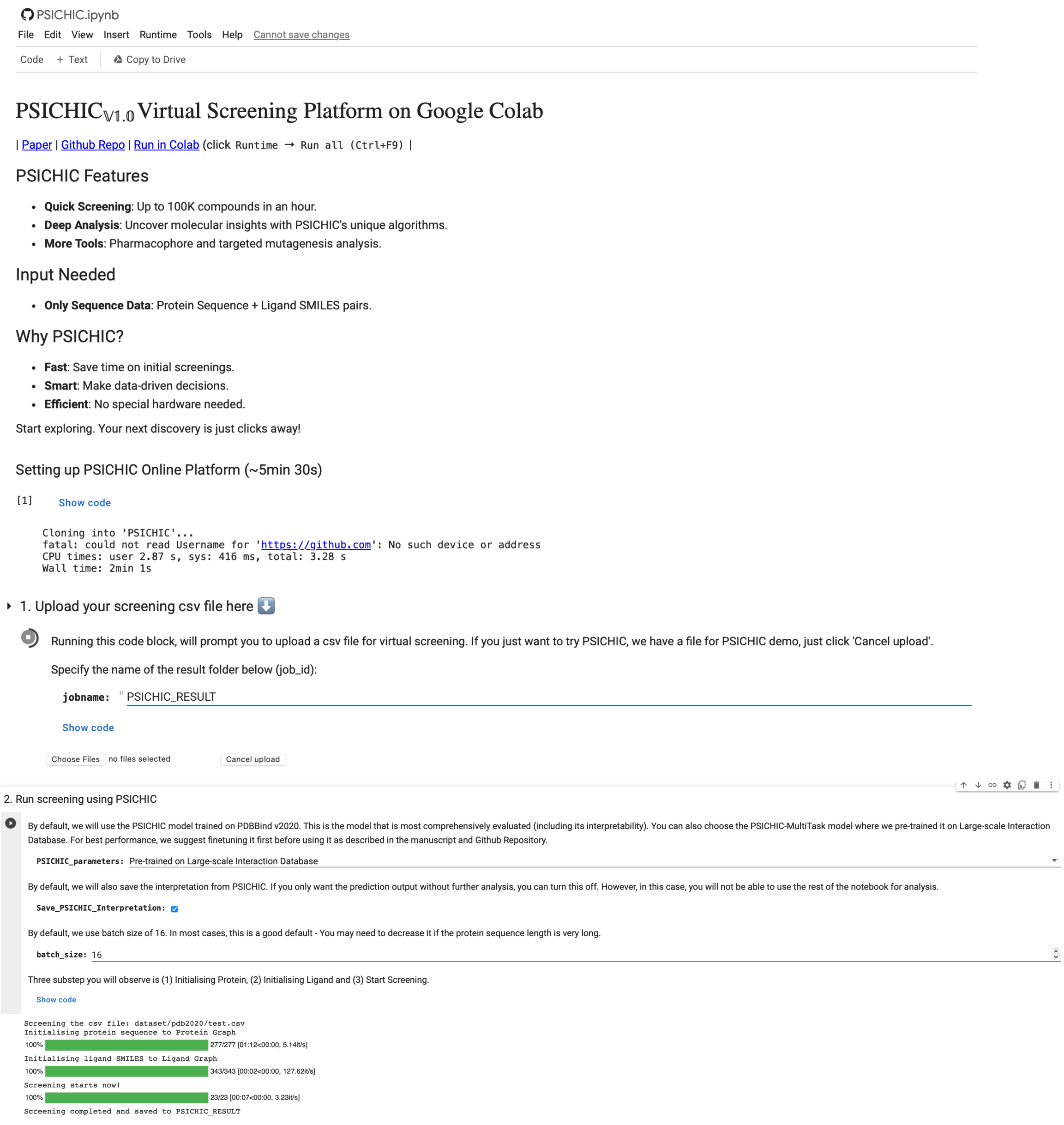

**Supplementary Fig. 1. PSICHIC Online Platform: Sequence Data is All You Need.** The platform operates on Google Colaboratory, offering accessibility without the need for specialized hardware. Users are required to input a CSV file containing Protein Sequence and Ligand SMILES strings, eliminating the need for high-resolution structural data.

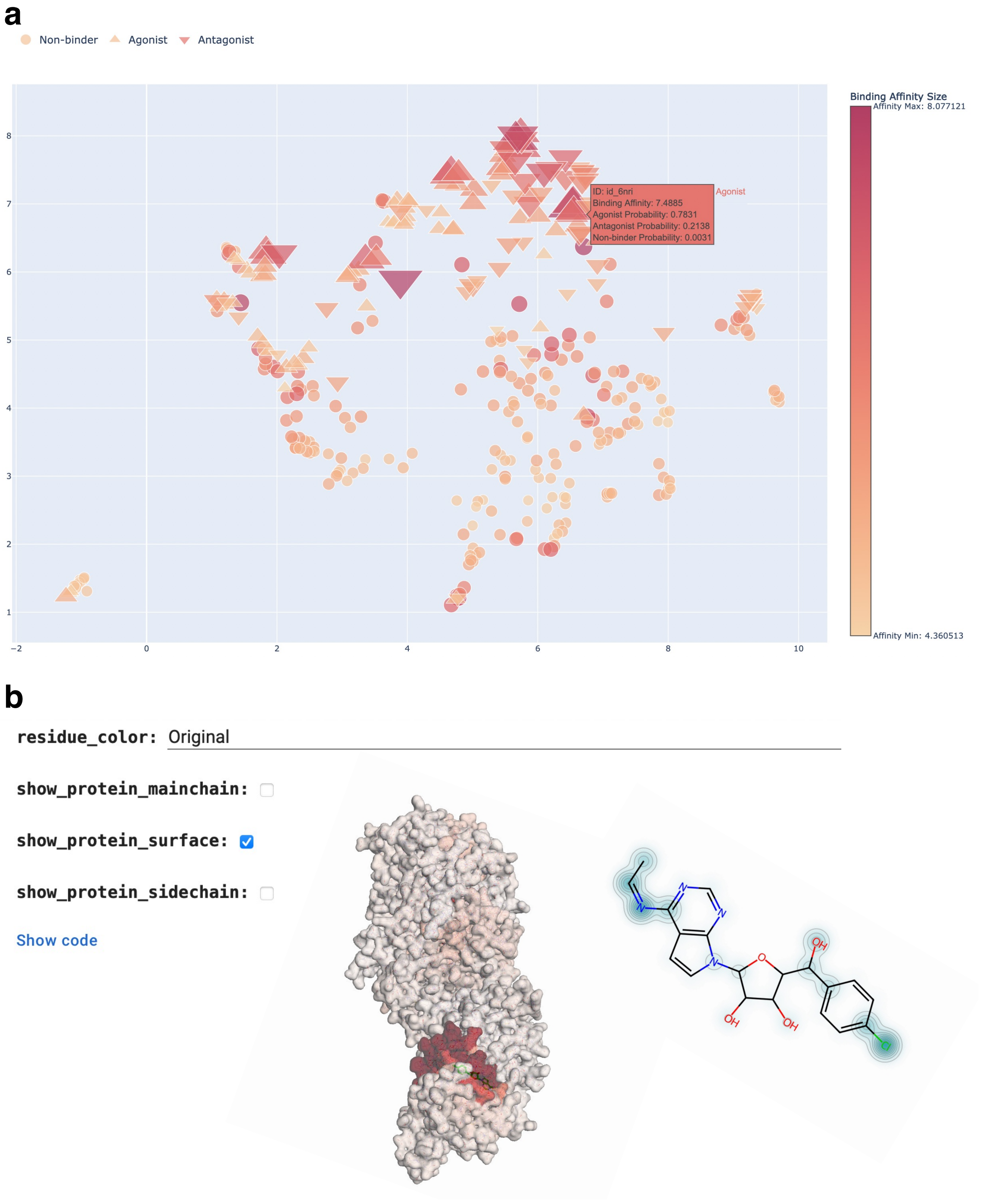

**Supplementary Fig. 2. PSICHIC Online Platform: Interactive Drug Discovery.** The PSICHIC online platform offers an interactive interface for drug discovery, allowing users to delve into multi-faceted analyses of protein-ligand interactions. It predicts both binding affinity and functional effects, and provides interpretable fingerprints that identify key residues in the protein and atoms in the ligand involved in the interaction. This interactive feature enhances the platform's utility by offering real-time insights into the underlying mechanisms of interaction.
